# Climate explains population divergence in drought-induced plasticity of functional traits and gene expression in a South African *Protea*

**DOI:** 10.1101/478230

**Authors:** Melis Akman, Jane E. Carlson, Andrew M. Latimer

**Author notes:** Corresponding author: Melis Akman.

## Abstract

Long term environmental variation often drives local adaptation and leads to trait differentiation across populations. Additionally, when traits change in an environment-dependent way through phenotypic plasticity, the genetic variation underlying plasticity will also be under selection. These processes could create a landscape of differentiation across populations in traits and their plasticity. Here, we studied drought responses in seedlings of a shrub species from the Cape Floristic Region, the common sugarbush (*Protea repens*). We measured morphological and physiological traits, and sequenced whole transcriptomes in 8 populations that represent both the climatic and the geographic distribution of this species. We found that there is substantial variation in how populations respond to drought, but we also observed common patterns such as reduced leaf size and leaf thickness, and upregulation of stress- and down-regulation of growth-related gene groups. Both high environmental heterogeneity and milder source site climates were associated with higher plasticity in various traits and co-expression gene networks. Associations between traits, trait plasticity, gene networks and the source site climate suggests that temperature may play a bigger role in shaping these patterns when compared to precipitation, in line with recent changes in the region due to climate change. We also found that traits respond to climatic variation in an environment dependent manner: some associations between traits and climate were apparent only under certain growing conditions. Together, our results uncover common responses of *P. repens* populations to drought, and climatic drivers of population differentiation in functional traits, gene expression and their plasticity.

## Introduction

Populations can persist in changing environmental conditions through three mechanisms: by tracking their climatic niche in space through migration or dispersal, by shifting their niche through adaptation, and/or by matching traits to the new environment through phenotypic plasticity (Chevin, Lande, and Mace 2010). For plants, plasticity can be the fastest response to environmental change, since it involves rapid adjustments of morphology, physiology and gene expression within a single generation. Like mean trait values, the form and magnitude of trait plasticity can be under selection and thus plasticity can also vary among populations along environmental gradients (Geber and Griffen 2003; Wennersten and Forsman 2012; Lázaro-Nogal et al. 2016, 2015; Wadgymar, Mactavish, and Anderson 2018). Previous studies have demonstrated that such population differences in plasticity might or might not have climatic drivers (Pratt and Mooney 2013; Lázaro-Nogal et al. 2015; Pohlman, Nicotra, and Murray 2005; Cochrane et al. 2015).

While plasticity can have strong consequences on functional traits (e.g. (Etterson 2004), it remains less clear how often plasticity confers environment-specific fitness benefits (Forsman 2015; Acasuso-Rivero et al. 2019). Plasticity has been directly shown to be adaptive: for example, Dudley and Schmitt (1996) showed that plastic stem elongation was beneficial at high but not low plant densities. Other studies have characterized patterns of plasticity that suggest adaptive plasticity (Sultan 2003; Nicotra et al. 2015; Baythavong 2011; Lázaro-Nogal et al. 2016; Cavender-Bares and Ramírez-Valiente 2017). Resurrection studies using historical and modern *Daphnia magna* populations showed evolution of plasticity for heat tolerance and fish predation (Cuenca Cambronero et al. 2018; Stoks et al. 2016). In light of these studies and theoretical predictions (Grime 1994; Bradshaw and Hardwick 1989; Alpert and Simms 2002; Schlichting and Pigliucci 1995; West-Eberhard 2003; Ghalambor et al. 2015), we expect that selection should promote local adaptation in plasticity under two main circumstances. First, plants from environments with strong and relatively predictable forms of spatial or temporal heterogeneity should be more plastic than those from homogeneous environments (Ackerly et al. 2000; Van Kleunen and Fischer 2005; Baythavong and Stanton 2010; Scheiner 2013; Lázaro-Nogal et al. 2015; Bonamour et al. 2019; Chevin and Hoffmann 2017; Reed et al. 2010); but see Palacio-López et al. (2015). Second, plants from harsh climates are expected to be less plastic than those from milder climates (Vitasse et al. 2013; Gugger et al. 2015), because the fitness costs of mismatch between the phenotype and the environment should be greater in resource-poor conditions (Alpert and Simms 2002). Elevation contrasts provide one example of this latter pattern: for an alpine herb, *Wahlenbergia ceracea*, inhabiting high montane and alpine elevations, plants from colder, higher-elevation environments tend to have both lower and less plastic growth rates than plants from warmer sites (Nicotra et al. 2015).

The form and magnitude of functional plasticity which organisms express can be expected to influence the vulnerability of populations to environmental change. This study focuses on a widespread plant species from the Cape Floristic Region (CFR) of South Africa, a biodiversity hotspot (Cowling 2009), that is predicted to be strongly affected by climate change. Our study species, *Protea repens* (common sugar bush), is the most widespread species in this plant genus with 112 species, ~85% of which are endemic (Sauquet et al. 2009; Rebelo 2001). Climate models project warming up to 3 °C by 2070 (Wilson, Latimer, and Silander 2015), and precipitation projections being more variable, most models project drying in the super-diverse western mountains (Wilson, Latimer, and Silander 2015; Hewitson and Crane 2006). Most Mediterranean-climate regions, including California and Southern Australia as well as the CFR, have recently experienced unprecedented droughts, which caused large mortality events in trees (Young et al. 2017; Allen et al. 2010) and shrubs (Valliere et al. 2017). Thus, understanding the patterns and effects of drought plasticity in functional traits is a vital issue for plant vulnerability in these semi-arid regions.

Plasticity in key plant traits, including stomatal density and conductance, growth rate, and leaf morphology, has been observed to vary among populations in long-lived woody plants (e.g. (Aubin et al. 2016) as well as herbaceous annuals (Jacobs and Latimer 2012). Likewise, along the environmental gradients *Protea* species inhabit, we have found evidence for local adaptation and environment-dependent benefits in several traits *(Carlson, Adams, and Holsinger 2016; Carlson, Holsinger, and Prunier 2011; Latimer et al. 2009; Mitchell, Carlson, and Holsinger 2018; Prunier, Holsinger, and Carlson 2012)*. Specifically, we found evidence for positive selection on stomatal density in populations from hotter and drier sites, with links to higher stomatal conductance in *P. repens (Carlson, Adams, and Holsinger 2016)*. Additionally, in the monophyletic group of white proteas, *P. subvestita* shows correlations between root length and rainfall seasonality. Biomass also decreases with increasing elevation in *P. mundii*, *P. punctata* and *P. lacticolor*, but shows the opposite trend in *P. mundii* (Prunier, Holsinger, and Carlson 2012). On a broader phylogenetic scale, we also found associations between temperature and plant size related traits such as leaf area and plant height (Mitchell, Carlson, and Holsinger 2018). There is also evidence for differential developmental plasticity and drought plasticity in leaf traits of several other *Protea* species (*Protea sect. Exsertae*: (Carlson and Holsinger 2012; Heschel et al. 2014).

Previously, we also showed that gene expression in populations grown in a common garden show patterns of differentiation corresponding to climatic variables such as elevation (a good proxy for temperature in CFR), minimum winter temperature and mean annual precipitation in *P. repens* (Akman et al. 2016). These gene networks also show correlations with key traits such as height, leaf area, stomatal density/size, providing insights into the genes involved in climate driven differentiation. Measurements of plasticity in gene expression, in which thousands of “phenotypes” can be measured simultaneously, are also valuable for investigating the causes and consequences of plasticity. Gene expression patterns can change within seconds of a stimulus, long before larger biochemical shifts and morphological changes become apparent. Gene expression patterns can also be linked to morphological and physiological adjustments, potentially providing insights into the functional genetic basis of adaptation and/or plasticity (Whitehead 2012; Akman et al. 2016) We note that such links do not pinpoint the genetic locus associated with the phenotypic variation, as loci influencing expression may lie in regulatory regions or in others genes whose expression is upstream of the focal gene. Previous studies have revealed patterns of adaptive plasticity in gene expression in fish (Dayan, Crawford, and Oleksiak 2015; Morris et al. 2014; Schneider et al. 2014), plants (Lasky et al. 2014; Colicchio et al. 2015) and corals (Barshis et al. 2013; Granados-Cifuentes et al. 2013). These studies demonstrate that gene expression profiling can usefully complement standard functional trait measurements to help understand the functional genetic basis and physiological significance of plasticity and how it varies across populations.

With this study, we aim to answer two questions: 1) How do populations of *P. repens* respond to drought, both in terms of functional traits and gene expression, together with their plasticity, and how do these responses vary across populations? and 2) Do the climatic conditions of the source sites covary with the mean values and plasticity of functional traits and gene expression levels? To do so, we imposed a severe, experimental drought on seedlings in a dry down experiment, and measured functional traits and gene expression patterns. Then, we tested for population differentiation in trait means and drought-response plasticity in functional traits and gene expression patterns, and evaluated associations between patterns of population differentiation and their potential climatic drivers.

## Materials and Methods

### Species description

*Protea repens* L. is a common evergreen shrub endemic to South Africa, with particularly high abundance in semi-arid fynbos of the CFR (Rebelo 2001). This species has narrow, sclerophyllous leaves and reaches 2-4 m in height. All viable and nonviable fruits remain within a woody, serotinous infructescence until the adult plant is killed by fire, which occurs at ~10-20 year intervals (with variation among geographic regions (Van Wilgen, Forsyth, and De Klerk 2010). Fruits disperse by tumbling, and seeds germinate during the next rainy season (May-Aug for much of the CFR). Due to the Mediterranean rainfall patterns in the western CFR, some western sites receive far more wintertime precipitation than do eastern population sites.

### Field sampling and climate data

In 2011, we collected seeds from 8 wild populations of *P. repens* spanning its geographic distribution in the CFR (**Figure 1A** and **Table S1**). The populations encompass a range of precipitation levels and degrees of seasonal concentration -- some sites have strong winter rainfall seasonality, while others experience both winter precipitation and some summer monsoonal rain (Table S1). The elevation range was deliberately wide, from ~150m to ~1500m. The reason why the lower elevation, warmer sites are “milder” is that they have little or no frost in winter, while being only moderately dry. High-elevation sites, in contrast, receive snow in winter and have high winds and shorter growing seasons. There are hot, dry, low-elevation sites in the interior of the Cape region that are not mild, but these tend to be too dry to support our study species, and were not included in our sampling. From 40-50 plants per population (mean n= 45 plants), we collected 1-2 mature seed heads from the previous year’s growth. Plants were sampled along transects through the population centre, with at least 1 m between each sampled plant for small populations (<100 plants), and 5 m for larger populations. Seed heads were allowed to air-dry until achenes were released. GPS coordinates of each site near the population center were used to extract elevation and six climate metrics for rainfall and temperature from GIS layers. A principal component analysis performed on these data confirmed that the sample sites cover most of the range of variation evenly in major environmental factors (**Figure 1B**).

**Fig 1.**
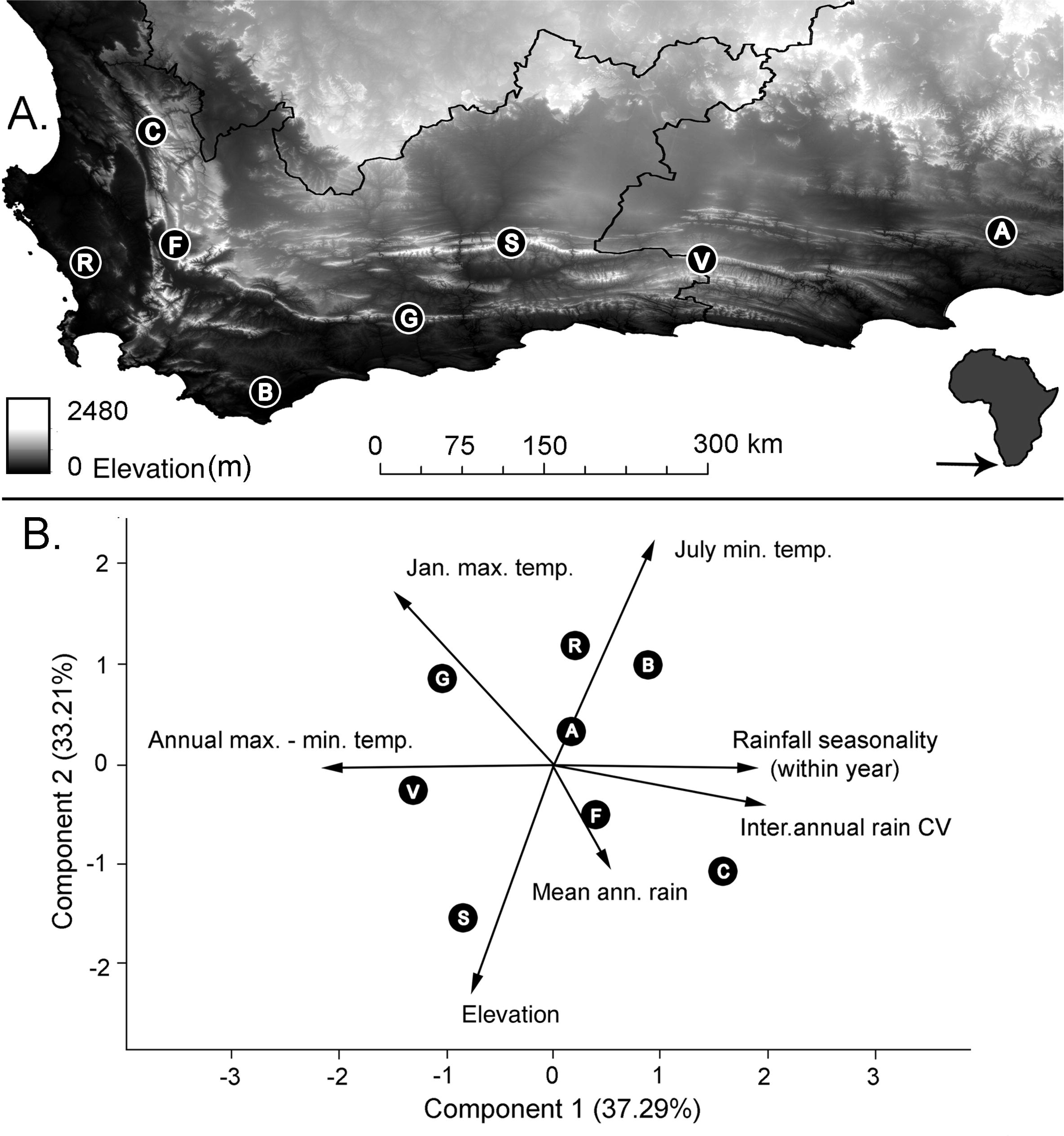
Map of CFR and locations of sampling sites (A); Principal component analysis for the climatic variables used in this study

The climate data layers were from the South African Atlas of Agrohydrology and Climatology (Roland Edgar Schulze 1997) with higher resolution for 4 and 5 below, and (R. E. Schulze 2007) for the rest, and were based on 50 year (collected between 1950-1999 when data was available from field stations) climate averages, extremes, or variances. The climate layers used were (1) mean annual rainfall, (2) average daily maximum temperature for summer (January), (3) average daily minimum temperature for winter (July), (4) precipitation concentration (an seasonality indicator we refer to as “rainfall seasonality”), measured as Markham’s concentration index (Markham 1970), which ranges from 0 to 100, with a value of 0 indicating perfectly even precipitation across all months, and 100 indicating that all precipitation falls in a single month (5) inter-annual rainfall variability, measured as the coefficient of variation in mean annual rainfall across years, and (6) extreme temperature range within a year, measured as the difference between summer maximum and winter minimum temperatures. Our seventh variable was elevation, which was extracted separately from ASTER GDEM (Daac 2011). Elevation is correlated strongly with winter minimum temperature, and we interpret it primarily as an indicator of temperature regime, although some other factors such as precipitation and wind speed are more weakly associated with elevation in the region (**Table S2**). Most of these layers have been used in our earlier work on *P. repens*, except for (5) and (6), which were added to complement rainfall seasonality as additional measures of environmental heterogeneity. Soil analyses for these source population locations were not available, but we standardized the general soil type: populations occurred (as is typical for this species) on low-nutrient, coarse, sandy soil derived primarily from quartzite sandstone parent material.

Only two pairs of climate variables were strongly correlated with each other: winter minimum temperature with elevation (Pearson’s R= −0.91) and summer maximum temperature with annual temperature range (R=0.70; **Table S2**). Rainfall seasonality in our dataset is positively associated with wintertime rainfall (total for Dec-Feb, Pearson’s R=0.80) and negatively associated with summertime rainfall (total for June-August, Pearson’s R=−0.90). The 12-year gap between the last collection year of the climate variables and seed collection date might hinder some of the associations we detect in later sections. However, the age of the plants (9-23 years based on veld age through burn history when available) from which seeds were collected will potentially eliminate this gap. We predict that selection should have already acted on the maternal plants and pollen sources, and their adaptations, if any, should be reflected in the germplasm we performed our experiments on.

### Experimental setup and dry-down treatments

In July 2014, we sowed seeds from 8 *Protea repens* populations on low nutrient mix (50% bark, 50% sand) in trays of 100 individual plugs 2×2 cm in size with a complete random design. Trays were placed in a long-day growth chamber (8 hours in dark at 8°C, 16 hours in light at 20°C). The trays were randomized every 3 days with each watering in the growth chambers. After 1.5 months, we transplanted germinated seedlings into small, deep pots (5 cm diameter x 17 cm deep) in trays of 50 each, in the same low nutrient mix. After transplanting, seedlings were rested for two weeks indoors under growth lights before being transferred to a single greenhouse bench at Nicholls State University Farm, Thibodaux, LA.

Two weeks post-greenhouse transfer and two weeks prior to the start of the dry-down experiment, we selected a total of 316 out of 419 seedlings (19-20 seedlings/population x 8 populations x 2 treatments) for uniform size for the experiment. We then reorganized the 2.5 month-old seedlings into trays containing 20-25 seedlings each, in a complete random design. In the experiment, seedlings were assigned to one of two treatments: control, in which we applied regular watering (~every 3 days, equal amounts of water was given to each plant) and drought, in which we stopped watering entirely for 12 days. Because we had a variable number of seedlings per maternal plant, we could not perfectly balance maternal lines across populations and treatments. To the extent possible, however, we included at least one seedling per maternal family per treatment; for 82% of the 83 maternal families, we were able to include at least one seedling per maternal family in each treatment, and for 51% of families, we included multiple seedlings per treatment (Table S3). The drought experiment began in late October 2014, when greenhouse conditions were warm but relatively dry, and thus most similar to summer conditions in the native range.

### Trait measurements and gene expression sampling

At 6 days and 12 days into the drought experiment, plants were measured for functional traits and leaves were sampled for transcriptomics. Our sampling design is depicted in **Figure S1**. Functional traits that were measured at both timepoints included plant height (cm), stomatal conductance, leaf area, leaf length:width ratio (LWR), specific leaf area (fresh leaf area/dry leaf mass; SLA), stomatal density (number per mm^2^), stomatal pore length (mm) and stem pigmentation. Stem pigmentation, measured as stem hue, was derived using Endler’s segment classification (Maia *et al.* 2013), based on the average of two reflectance spectra readings (400-700 nm) taken at mid-stem (JAZ, Ocean Optics). On the segment classification scale for hue, higher values represent redder stems that contain higher levels of anthocyanin. Stomatal conductance was measured using a steady-state leaf porometer (SC-1, Decagon) on one leaf per plant, with an additional adjacent leaf measured simultaneously if the chamber was not filled by a single leaf due to small leaf size.

All other leaf measurements were taken from the 2nd or 3rd youngest expanding leaf from the stem apex. Immediately following conductance measurement, the leaf was removed to measure leaf area, width, length and stomatal traits. Fresh leaves were digitally scanned shortly after harvest at 400 dpi on a flatbed scanner. After scanning, acrylic peels were taken from the adaxial leaf surface and placed on cellophane, which were later viewed under a light microscope at 40x for stomatal measurements (see Carlson *et al.* 2016). Although height and pigmentation were measured on all plants on both timepoints (mean sample size per population=39), all other traits were measured on only a subset of plants on each of these days, typically in association with the transcriptomic data (10-20 individuals per population for each day). Two additional variables were calculated by subtracting day 6 from day 12 values on the same plant: pigment accumulation (day 12 hue - day 6 hue) and growth (day 12 height - day 6 height). A principle component analysis was performed on the traits measured separately for day 6 and day 12. Details of sampling for gene expression and carbohydrate analysis are included in Supplementary Materials and Methods. Briefly, leaves were sampled at both time points for whole transcriptome sequencing, and at day 12, the whole plants were freeze dried and extracted with 70% methanol, and carbohydrates were measured with anthrone method (as described in (Akman et al. 2012)).

### Gene expression analyses and bioinformatics

See Supplementary Materials and Methods for transcriptome library preparation and sequencing details. Raw reads were trimmed and demultiplexed with trimmomatic (Bolger, Lohse, and Usadel 2014) and fastx toolkit (Hannon, n.d.). By using bowtie (Langmead and Salzberg 2012), trimmed reads were mapped to a previously published de-novo transcriptome of *P. repens* (Akman et al. 2016a) assembled by using sequences of individuals that also include all of our focal populations. Before reads were mapped to the transcriptome, a further clustering of the contigs were done using cd-hit (Fu et al. 2012). Read counts, extracted using eXpress (Pachter 2011), averaged 6.02 million reads per sample (ranged between 2.46-21.80 million). Contigs with <1 counts per million (cpm) were excluded from the analyses. Differential gene expression analyses were performed using edgeR. For each time point, differentially expressed genes per population were revealed using pairwise comparisons using negative binomial models. The common set of differentially expressed genes was obtained by extracting the overlap of differentially expressed genes for each population at day 12 (**Table S5**). We then used linear models in edgeR, including sequencing lane as a fixed factor, to detect contigs that showed variation between treatments and among populations (population x treatment). Multi-dimensional scaling (MDS) analysis was performed for each time point to visualize general patterns of gene expression for all the samples using the top 500 genes that explain the most variation (**Figure 2A-B and Figure S6**).

**Fig 2.**
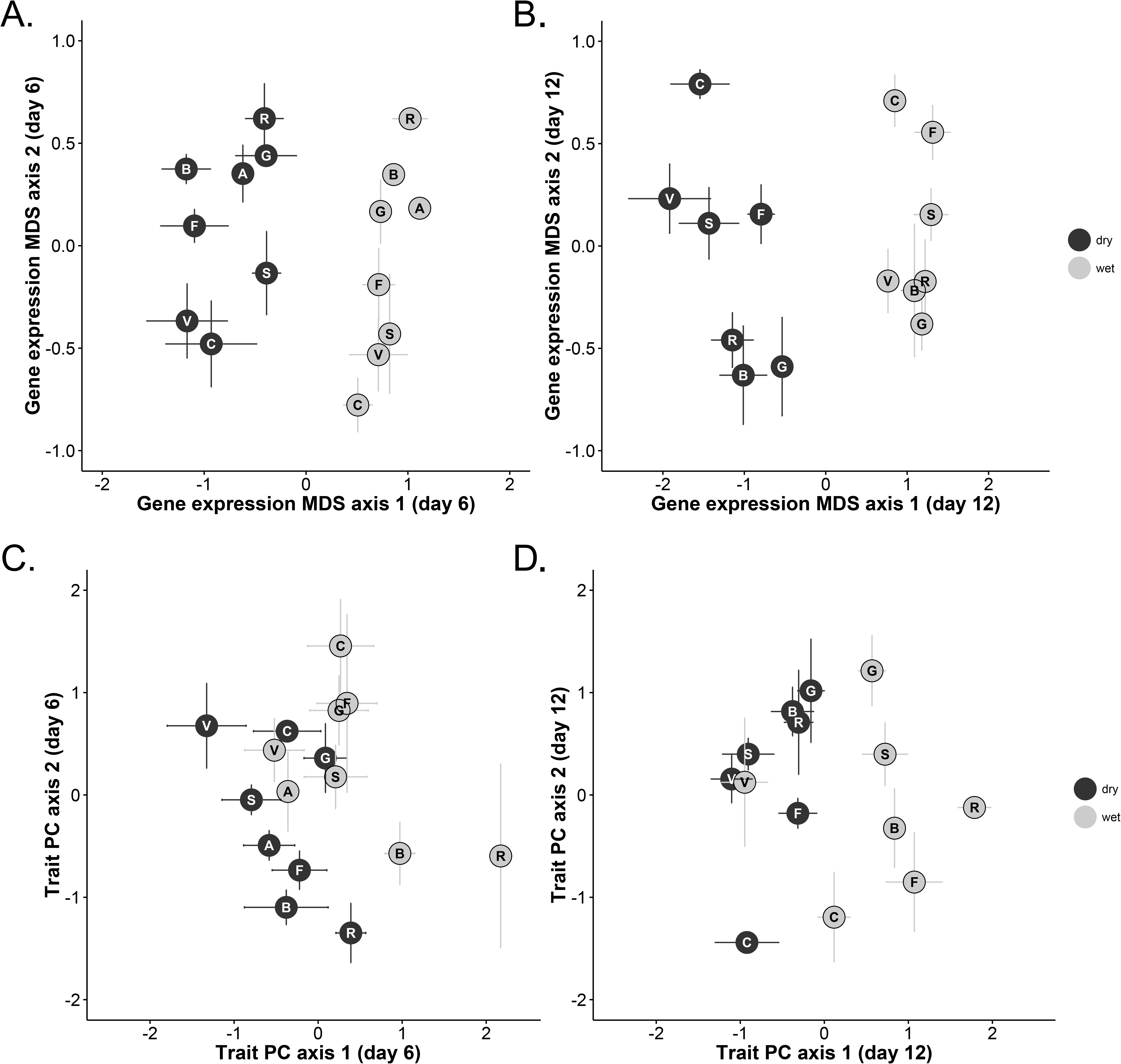
Multi-dimensional scaling (MDS) plots for gene expression patterns at day 6 (A) and day 12 (B); principal component analysis of measured traits at day 6 (C) and day 12 (D) into the drought treatment.

Co-expression gene networks were computed using WGCNA (Langfelder and Horvath 2008) with consensus clustering option for the 2 timepoints together. Normalized and cpm-filtered read counts were transformed with “voom” in limma package. A soft-threshold of 6 was used for clustering and 14 co-expressed gene networks were constructed. The number of genes within each gene network varied between 49 (GN13) to 7682 (GN1). The gene network “grey” (called GN0 in this study) as constructed by WGCNA for genes that do not fit any gene network has 3124 genes. A representative “eigen gene” value, which is the first principal component axis for all genes within the network, was used for further analyses of climate and trait associations. By using gene orthology of our genes with Arabidopsis (Akman et al. 2016a), we also tested for Gene Ontology (GO) enrichment for the 13 networks. This analysis was performed in Gorilla (Eden et al. 2009), using the network genes as target set and the whole gene set as the background. Please note that, when we refer to a statistical test for a given gene network (GN), an association or a linear mixed model test, we are referring to the eigengene value of that gene network. However, when we mention an associated function, this is based on all the genes on that network and the corresponding GO analyses. Contigs within each module, GO enrichments and the genes with the highest module membership for a single network are provided in **Table S4**.

### Statistical analyses

#### How do populations respond to drought and how do these responses vary?

We tested whether drought significantly affected traits in *Protea repens* using linear mixed models. We used a total of 23 plant metrics in our analyses, including 8 traits on day 6, and 15 traits on day 12. We analyzed each trait on each day separately, because preliminary analyses on combining both days revealed significant timepoint effects for almost every response variable. Each model included the fixed effects of treatment, population, and the population by treatment interaction (G x E) and random effects of maternal line nested within population. In these models, the treatment-by-population interaction term indicates whether there was differential plasticity among populations, or in other words, that plants from different populations responded differently to the treatment (G x E effect). These analyses were performed using the lme4 package (Bates, Maechler, and Bolker 2012) in R 3.1 (Team 2014). To obtain p-values for each fixed effect, we successively dropped one predictor from the full model and used a chi-square test to compare AIC scores between models. Given the large number of statistical tests that this entailed, we also adjusted p-values according to estimated false discovery rates (FDR; R package “fdrtool”, local method).

Gene expression analyses followed the same general approach as for traits. As response variables, these analyses focused on eigengenes in the 14 gene networks (GN) that were produced through the gene clustering analysis. Models for gene networks included fixed effects of treatment, population, and their interaction, random effects of maternal line nested within population, and a random effect of sequencing lane. We also tested for genes that are differentially expressed within a population. Commonly regulated genes for all populations are given in **Table S5**. We also tested for genes differentially expressed among populations for both timepoints and days separately.

#### Do the climatic conditions of the source sites covary with traits, gene expression and their plasticity?

To test whether characteristics of the home climate predicted population variation in traits and gene expression, we used mixed effect models. In these models, traits and gene networks (GN) were response variables, the climate variables were fixed effect terms, and maternal line nested in population was a random effect variable. For testing the significance of the fixed factors, we calculated p-values based on chi-square tests as explained above. FDR corrections were applied to p-values in two groups: traits and gene networks. Day 6 and Day 12 FDR corrections were also applied separately. For variables with significant climate and treatment interaction, each treatment was also analyzed separately to estimate slopes and test their significance within treatments.

We used another set of linear mixed models to test whether there were significant associations between traits and gene expression patterns. We performed these analyses with lme4 in R as above, with each pairwise comparison between a trait and a gene network in its own model. Response variables were one of 8 (at day 6) or 15 (at day 12) traits and predictor variables were one of 14 eigen gene values for a gene network. Random effects were sequencing lane, population and maternal line nested within population. Correlations were tested separately for the two timepoints, and p-values were calculated as explained previously. An FDR correction was applied separately to the two time points (Table S6).

To characterize plasticity for individual traits and to quantify the direction and magnitude of plasticity, we used maternal line plasticity indices. For each maternal family, we calculated plasticity as the family mean trait value in the wet treatment minus the family mean trait value in the drought treatment, divided by the overall family mean trait value (for families with only one plant per treatment, we used the trait value of that plant rather than a family mean). Note that this plasticity metric can take on both positive and negative values: values close to 0 indicate low plasticity (little difference between treatments), while large negative and large positive values both indicate high plasticity.

We used mixed effect models to determine whether characteristics of the home climate predicted cross-population variation in plasticity of traits and gene expression. In our analyses, plasticity in traits and gene networks were response variables (these are the above mentioned family means), the 7 climate variables were fixed effect variables and population was included as a random effect. Each climate variable was tested against each response variable in a separate model, with day 6 and day 12 measurements analyzed separately (lmer library). In order to do this, we used a subset of gene network and trait pairs that already showed a climate x treatment effect (before FDR-correction;**Table S7&8**). These included 5 and 11 tests for gene networks, and 2 and 15 for traits for day 6 and day 12, respectively.

## Results

### How do responses to drought vary among populations?

#### Drought responses are stronger in gene expression compared to traits

Six days into the drought experiment, only three of the 8 traits differed significantly between treatments: unwatered plants had shorter stems, lower stomatal conductance and more variable SLA (**Table 1**). At the later stage, treatment effects became pronounced for more traits measured (12 of 15): unwatered plants had lower stomatal conductance, decreased height, slower growth in height (from day 6 to day 12), higher SLA, smaller leaf area, denser stomata, smaller stomatal pore size, increasing pigment accumulation, lower carbohydrates aboveground (but not belowground). In the earlier stages of the stress, morphological and physiological traits did not show responses as pronounced as those observed in gene expression patterns, potentially due to time required for morphological changes to happen (**Figure 2**). In contrast to most of the traits, the treatment effect was significant for over half of the gene networks already at day 6 into the treatment. This trend of more pronounced gene expression response was visible both in co-expression gene networks (**Table 3**, **Figure S2**) and Multi-dimensional Scaling (MDS) analyses (**Figure 2A-B**), which indicates a lag between the rapid adjustments in gene expression and trait onset. However, we should also note that the MDS for gene expression is based on the top 500 genes that explain the most variation, and we might not have measured morpho-physiological traits that might also respond as quickly. Notably, population V (Baviaanskloof, a high-elevation eastern site) did not show a treatment effect in traits at the later stage, although the effect was quite pronounced in gene expression (**Figure 2B&D** and **3A**).

**Table 1.** Results of linear mixed models testing trait differences of *Protea repens* seedlings at 6 and 12 days into the dry down experiment. The model compares values between watered and unwatered treatments as well as among 8 wild-sourced populations and the population by treatment interaction. Significant values are bolded.

| Response variable | Fixed effects | day 6 |  |  |  | day 12 |  |  |  |
| --- | --- | --- | --- | --- | --- | --- | --- | --- | --- |
|  |  | df | Chi-sq | P-value | q-val | df | Chi-sq | p-value | q-value |
| Leaf area (cm<sup>2</sup>) | Population | 7 | 49.91 | **<0.0001** | **<0.0001** | 6 | 69.16 | **<0.0001** | **<0.0001** |
|  | Treatment | 1 | 0.17 | 0.678 | 0.881 | 1 | 23.39 | **<0.0001** | **<0.0001** |
|  | Pop. × Trt. | 7 | 8.61 | 0.282 | 0.423 | 6 | 13.3 | 0.039 | 0.050 |
| SLA (cm<sup>2</sup>/g)<br>Log transform | Population | 7 | 5.15 | 0.642 | 0.881 | **6** | **15.35** | **0.018** | **0.027** |
|  | Treatment | **1** | **12.13** | **0.0005** | **0.002** | 1 | 73.14 | **<0.0001** | **<0.0001** |
|  | Pop. × Trt. | **7** | **19.23** | **0.008** | **0.027** | **6** | **16.71** | **0.010** | **0.017** |
| LWR | Population | 7 | 67.48 | **<0.0001** | **<0.0001** | 6 | 78.55 | **<0.0001** | **<0.0001** |
|  | Treatment | 1 | 0.06 | 0.814 | 0.881 | 1 | 1.86 | 0.173 | 0.210 |
|  | Pop. × Trt. | 7 | 3.47 | 0.839 | 0.881 | 6 | 1.52 | 0.958 | 0.958 |
| Stomatal density | Population | **7** | **17.46** | **0.015** | **0.040** | **6** | **29.73** | **<0.0001** | **0.0001** |
|  | Treatment | 1 | 3.07 | 0.080 | 0.192 | **1** | **12.95** | **0.0003** | **0.0007** |
|  | Pop. × Trt. | 7 | 9.56 | 0.215 | 0.369 | 6 | 3.59 | 0.732 | 0.765 |
| Stomatal pore size (um) | Population | 7 | 3.87 | 0.795 | 0.881 | **6** | **21.92** | **0.001** | **0.002** |
|  | Treatment | 1 | 0.04 | 0.844 | 0.881 | **1** | **27.68** | **<0.0001** | **<0.0001** |
|  | Pop. × Trt. | 7 | 3.05 | 0.881 | 0.881 | **6** | **14.27** | **0.027** | **0.038** |
| Stem pigmentation | Population | 7 | 34.13 | **<0.0001** | **<0.0001** | 7 | 38.51 | **<0.0001** | **<0.0001** |
|  | Treatment | 1 | 1.85 | 0.174 | 0.336 | 1 | 0.42 | 0.515 | 0.579 |
|  | Pop. × Trt. | 7 | 10.12 | 0.182 | 0.336 | 7 | 5.71 | 0.574 | 0.630 |
| Stem height (cm) | Population | **7** | **105.34** | **<0.0001** | **<0.0001** | 7 | 121.24 | **<0.0001** | **<0.0001** |
|  | Treatment | **1** | **5.9** | **0.015** | **0.040** | 1 | 49.4 | **<0.0001** | **<0.0001** |
|  | Pop. × Trt. | 7 | 4.1 | 0.768 | 0.881 | 7 | 6.86 | 0.443 | 0.511 |
| Stomatal conductance | Population | 7 | 0 | 0.254 | 0.406 | **6** | **17.52** | **0.008** | **0.014** |
|  | Treatment | **1** | **0** | **<0.0001** | **<0.0001** | **1** | **104.7** | **<0.0001** | **<0.0001** |
|  | Pop. × Trt. | 7 | 0 | 0.148 | 0.323 | **6** | **23.43** | **0.0007** | **0.001** |

At day 6 into the treatment, unwatered plants had higher eigengene values for gene networks GN 2, 4, 5, 6, and 7 and lower values for gene networks GN 1, 9 and 11, relative to watered plants (**Table 3**, **Figure S3**). These modules show gene enrichment in GO categories for ATPase activity (GN2), oxidoreductase activity (GN4), cytoskeleton organization (GN5), photorespiration (GN6), response to high light intensity and temperature (GN7; note that p-value<0.0005, fdr-corrected p-value is not significant), cell wall organization (GN1) and pyridine-containing compound metabolic process (GN9). As the stress advanced after 12 days into the drought treatment, watered and unwatered plants differed significantly for most traits and gene networks. Significant treatment responses were detected for 10 of 14 gene networks and 12 of 15 traits. (**Tables 1–3**, **Figures S3-5**). In addition to the gene networks with treatment effects at day 6, GN3 (tropism and carbohydrate metabolism related, and shows associations with leaf area, height and aboveground soluble carbohydrates) and GN13 (homeostasis related) also had evident treatments effects at day 12, with GN3 having lower values and GN13 higher values under drought.

#### Drought up-regulates stress related genes and down-regulates growth related genes

The number of genes differentially regulated between earlier and later stages of the stress and among populations differed significantly (**Figure 3A&B**). For example, population G (Garcia’s Pass) does not have any genes differentially expressed at day 6, indicating either a lower level of stress or lack of accurate perception and timely signal transduction of the environmental cues to gene expression phenotypes. In contrast, population B (Bredasdorp) had several hundred differentially expressed genes by day 6. All populations had at least several hundred differentially expressed genes by day 12. Two populations (C-Cederberg and S-Swartberg) showed few differentially expressed genes initially, but over 1500 differentially expressed genes by day 12 (**Figure 3A**). Although the number of genes regulated upon drought stress was highly variable, we found 110 and 63 genes to be consistently up-or down-regulated, respectively, among populations (**Table S5**). The up-regulated genes include heat shock proteins, auxin signaling and ABA catabolism genes, genes related to drought stress and photosynthesis, and a leaf amylase; whereas down-regulated genes included ARF genes (involved in cell elongation and division), and salt stress-, fatty acid and amino acid biosynthesis-related genes.

**Fig 3.**
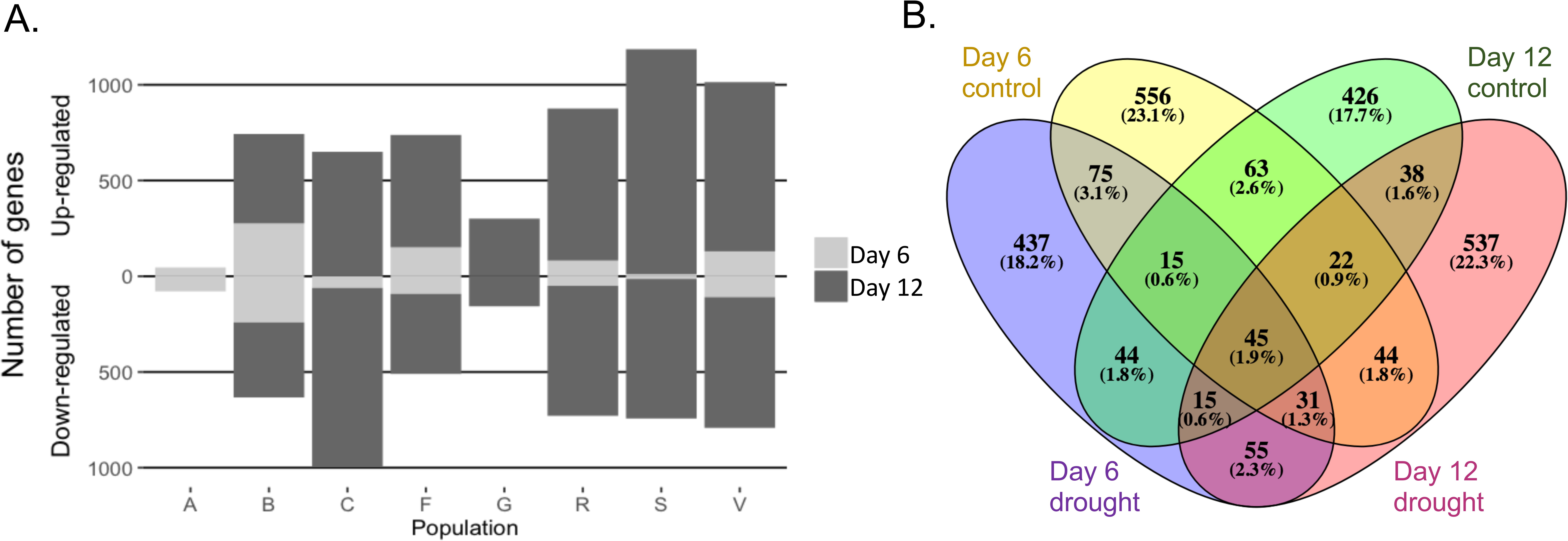
Number of differentially expressed genes for the 8 populations for after 6 and 12 days of the drought treatment (A). Note that population A is not sampled at day 12, and population G does not have any differentially expressed genes between the treatments at day 6. Number of differentially expressed genes among populations under control and drought conditions for day 6 and 12 separately (B). For example, the intersecting ellipse between Day6 control and DAY6 drought indicates number of genes that are differentially expressed among populations, regardless of the treatment.

#### Population differentiation in stress response becomes more evident as the stress advances

Variation among populations is evident in most of the traits, 11 out of 15 traits were significantly different between the populations at day 6, and all traits except for root length showed a population effect at day 12 (**Table 1&2**). We also observed G x E effects (variation in population responses to drought) to a limited extent on day 6, and to a greater extent on day 12 (**Table 1**). At day 6, only SLA and gene network 13 (enriched for homeostasis GO term) showed variation in responses to drought across populations (**Table 1&3**). SLA was lower under drought for western populations (R, G, C and F), but it was higher under drought in the remaining 4 populations (V, A, B, S; **Table 1**, **Figure S3**). At day 12, populations showed variation in their response to drought for SLA, stomatal size, pigment accumulation, stomatal conductance, stem growth rate, starch biomass aboveground, sugar biomass above and belowground. In addition to GN13 at day 6, GN7 expression (linked to high light intensity and high temperature GO term) also showed a G x E effect at day 12 (**Table 3**).

**Table 2.** Results of linear mixed models testing for differences in growth and carbohydrate storage between watered and unwatered *Protea repens* seedlings and among 8 populations at the end of the dry-down experiment (day 12). Significant values are bolded.

| Response variable | Fixed effects | df | Chi-sq | p-val | q-val |
| --- | --- | --- | --- | --- | --- |
| Soluble sugars mass (g) belowground | Population | **7** | **55.57** | **<0.0001** | **<0.0001** |
|  | Treatment | **1** | **6.33** | **0.012** | **0.019** |
|  | Pop. × Trt. | **7** | **17.52** | **0.014** | **0.022** |
| Soluble sugars mass (g) aboveground | Population | **7** | **80.08** | **<0.0001** | **<0.0001** |
|  | Treatment | **1** | **25.69** | **<0.0001** | **<0.0001** |
|  | Pop. × Trt. | **7** | **24.27** | **0.001** | **0.002** |
| Starch mass (g) belowground | Population | **7** | **21.06** | **0.004** | **0.008** |
|  | Treatment | 1 | 0.10 | 0.748 | 0.765 |
|  | Pop. × Trt. | 7 | 4.57 | 0.712 | 0.763 |
| Starch mass (g) aboveground | Population | **7** | **57.40** | **<0.0001** | **<0.0001** |
|  | Treatment | **1** | **60.10** | **<0.0001** | **<0.0001** |
|  | Pop. × Trt. | **7** | **100.76** | **<0.0001** | **<0.0001** |
| Growth rate (height on Day 12-6) | Population | **7** | **34.85** | **<0.0001** | **<0.0001** |
|  | Treatment | **1** | **194.34** | **<0.0001** | **<0.0001** |
|  | Pop. × Trt. | **7** | **26.33** | **0.0004** | **0.0009** |
| Root length (mm) | Population | 6 | 9.36 | 0.154 | 0.192 |
|  | Treatment | **1** | **6.66** | **0.010** | **0.017** |
|  | Pop. × Trt. | 6 | 6.3 | 0.391 | 0.463 |
| Stem pigment accumulation | Population | **7** | **15.47** | **0.031** | **0.041** |
|  | Treatment | **1** | **5.08** | **0.024** | **0.035** |
|  | Pop. × Trt. | **7** | **15.58** | **0.029** | **0.040** |

**Table 3.** Results of linear mixed models comparing eigengene values of the gene networks (GN0-GN13) at 6 and 12 days into the dry down experiment for watered and unwatered plants of *P. repens*. Significant values are bolded.

|  | Fixed effects | Fixed effects at 6 days |  |  |  | Fixed effects at 12 days |  |  |  |
| --- | --- | --- | --- | --- | --- | --- | --- | --- | --- |
|  |  | df | Chi-sq | p-value | q-val | ddf | Chi-sq | p-value | q-val |
| GN0 | Population | **7** | **24.89** | **0.0008** | **0.0042** | **6** | **80.38** | **<0.0001** | **<0.0001** |
|  | Treatment | 1 | 2.05 | 0.152 | 0.281 | 1 | 0.77 | 0.380 | 0.479 |
|  | Pop. × Trt. | 7 | 5.72 | 0.573 | 0.655 | 6 | 8.51 | 0.203 | 0.294 |
| GN1 | Population | **7** | **18.59** | **0.010** | **0.042** | **6** | **16.49** | **0.011** | **0.026** |
|  | Treatment | **1** | **91.40** | **<0.0001** | **<0.0001** | **1** | **123.77** | **<0.0001** | **<0.0001** |
|  | Pop. × Trt. | 7 | 11.30 | 0.126 | 0.252 | 6 | 6.71 | 0.348 | 0.457 |
| GN2 | Population | 7 | 8.91 | 0.259 | 0.363 | 6 | 13.87 | 0.031 | 0.0566 |
|  | Treatment | **1** | **13.01** | **0.0003** | **0.0018** | **1** | **50.11** | **<0.0001** | **<0.0001** |
|  | Pop. × Trt. | 7 | 9.65 | 0.210 | 0.351 | 6 | 3.22 | 0.781 | 0.841 |
| GN3 | Population | 7 | 8.56 | 0.286 | 0.387 | **6** | **18.70** | **0.005** | **0.013** |
|  | Treatment | 1 | 0.07 | 0.796 | 0.836 | **1** | **9.88** | **0.00167** | **0.005** |
|  | Pop. × Trt. | 7 | 9.30 | 0.232 | 0.351 | 6 | 1.47 | 0.962 | 0.962 |
| GN4 | Population | 7 | 16.16 | 0.024 | 0.078 | **6** | **36.29** | **<0.0001** | **<0.0001** |
|  | Treatment | **1** | **7.52** | **0.006** | **0.028** | **1** | **75.50** | **<0.0001** | **<0.0001** |
|  | Pop. × Trt. | 7 | 6.04 | 0.536 | 0.643 | 6 | 3.80 | 0.704 | 0.799 |
| GN5 | Population | 7 | 12.91 | 0.074 | 0.177 | **6** | **26.58** | **0.0002** | **0.0007** |
|  | Treatment | **1** | **13.37** | **0.0003** | **0.0018** | **1** | **68.66** | **<0.0001** | **<0.0001** |
|  | Pop. × Trt. | 7 | 9.32 | 0.231 | 0.351 | 6 | 2.90 | 0.821 | 0.841 |
| GN6 | Population | 7 | 12.84 | 0.076 | 0.177 | 6 | 11.76 | 0.067 | 0.113 |
|  | Treatment | **1** | **56.14** | **<0.0001** | **<0.0001** | **1** | **107.13** | **<0.0001** | **<0.0001** |
|  | Pop. × Trt. | 7 | 9.00 | 0.253 | 0.363 | 6 | 7.72 | 0.260 | 0.352 |
| GN7 | Population | 7 | 15.34 | 0.032 | 0.096 | **6** | **19.99** | **0.003** | **0.008** |
|  | Treatment | **1** | **67.46** | **<0.0001** | **<0.0001** | **1** | **86.20** | **<0.0001** | **<0.0001** |
|  | Pop. × Trt. | 7 | 10.67 | 0.154 | 0.281 | **6** | **14.79** | **0.022** | **0.049** |
| GN8 | Population | 7 | 15.11 | 0.035 | 0.098 | 6 | 3.69 | 0.719 | 0.799 |
|  | Treatment | 1 | 0.65 | 0.420 | 0.540 | 1 | 1.49 | 0.222 | 0.311 |
|  | Pop. × Trt. | 7 | 9.96 | 0.191 | 0.334 | 6 | 4.10 | 0.663 | 0.796 |
| GN9 | Population | 7 | 12.56 | 0.084 | 0.186 | **6** | **23.63** | **0.0006** | **0.0019** |
|  | Treatment | **1** | **26.83** | **<0.0001** | **<0.0001** | **1** | **102.74** | **<0.0001** | **<0.0001** |
|  | Pop. × Trt. | 7 | 6.21 | 0.515 | 0.636 | 6 | 13.97 | 0.030 | 0.057 |
| GN10 | Population | 7 | 1.76 | 0.972 | 0.972 | 6 | 10.58 | 0.102 | 0.165 |
|  | Treatment | 1 | 0.04 | 0.840 | 0.860 | 1 | 0.13 | 0.723 | 0.799 |
|  | Pop. × Trt. | 7 | 5.46 | 0.604 | 0.655 | 6 | 14.12 | 0.028 | 0.056 |
| GN11 | Population | 7 | 13.34 | 0.064 | 0.168 | 6 | 12.34 | 0.055 | 0.096 |
|  | Treatment | **1** | **31.91** | **<0.0001** | **<0.0001** | **1** | **64.04** | **<0.0001** | **<0.0001** |
|  | Pop. × Trt. | 7 | 9.27 | 0.234 | 0.351 | 6 | 6.33 | 0.388 | 0.479 |
| GN12 | Population | 7 | 7.05 | 0.424 | 0.540 | 6 | 9.38 | 0.154 | 0.231 |
|  | Treatment | 1 | 2.75 | 0.097 | 0.204 | 1 | 0.06 | 0.808 | 0.841 |
|  | Pop. × Trt. | 7 | 5.53 | 0.596 | 0.655 | 6 | 9.41 | 0.152 | 0.231 |
| GN13 | Population | 7 | 16.84 | 0.018 | 0.063 | 6 | 14.14 | 0.028 | 0.056 |
|  | Treatment | 1 | 0.26 | 0.608 | 0.655 | **1** | **30.16** | **<0.0001** | **<0.0001** |
|  | Pop. × Trt. | **7** | **17.85** | **0.013** | **0.0496** | **6** | **16.57** | **0.011** | **0.026** |

For each treatment, we also tested for genes differentially expressed among populations (testing for population effect) and timepoints separately. This analysis reveals which genes are differentially expressed among populations, indicating population differentiation, within a treatment and a timepoint. With these analyses, we find 851 and 668 genes to be differentially regulated among populations for the control treatment, and 717 and 786 for drought treatment at day 6 and 12, respectively (FDR-corrected p-value<0.01). We also observed some overlap between these groups (**Figure 3B**); 45 of the genes being differentially expressed among populations in both of the treatments for both days. Interestingly, the majority of genes differentially expressed in a treatment did not overlap between the two timepoints and this holds true for both of the treatments (e.g. purple and pink in **Figure 3B**). Likewise, the genes differentially expressed in the different treatments did not show overlap within a single timepoint (e.g. purple and yellow in **Figure 3B**). These results show that together with a strong treatment effect, there is a pronounced timepoint effect in our experiment for gene expression (note that the plants were all harvested at the same time of the day but 6 days apart).

#### Most gene networks are associated with physio-morphological traits

Testing for associations between gene networks and traits, we showed that 12 of the 14 gene networks show correlations with at least one trait (Table S6). For example, GN 4 (oxidoreductase activity), 5 (cytoskeleton organization), 7 (response to high light intensity and temperature), 9 (pyridine-containing compound metabolic process) and 13 (homeostasis) all show correlations with plant height at day 6. At day 12, these same gene networks and an additional GN3 (tropism and carbohydrate metabolism-related) are correlated with plant height. GN13 also shows correlation with leaf area and stomatal conductance at day 6, and additionally SLA, stomatal size, aboveground starch and soluble carbohydrates at day 12. GN4 and 5 show correlations with SLA and leaf area at day 6, and above- and belowground soluble carbohydrates additionally at day 12. Similarly, GN6 (photorespiration) is associated with leaf area, stomatal conductance, aboveground starch and soluble carbohydrates and belowground soluble carbohydrates at day 12. Additionally, GN7 shows correlations with SLA at day 6, and GN12 (no GO enrichments) shows correlations with leaf area at day 12. GN9 shows correlations with leaf area and SLA at both time points, and with conductance at day 6 and above ground soluble carbohydrates and starch and below ground soluble carbohydrates at day 12.

### Do the climatic conditions covary with traits, gene expression and their plasticity?

#### Gene networks and functional traits correlate with temperature

In order to assess climatically driven trait and gene expression variation among populations, we tested for correlations between source-site climate variables and co-expression gene networks and functional traits. We found several gene networks that covary with climatic variables (**Figure 4**; boxed combinations). These include GN 3, 4, 5, 9 and 13 at day 6, and GN3, 4, 5, 6, 7, 10, 12 and 13 at day 12. Most of these gene networks showed strong correlations with elevation and the closely related variables winter minimum temperature and summer maximum temperature. We also observed correlations between mean annual precipitation and GN6, 10 and 13, stomatal conductance and starch content in shoots at day 12. Stem growth showed correlations with elevation, summer maximum and winter minimum temperature. Aboveground starch content co-varied with elevation, mean annual precipitation and intra-annual precipitation variation. Similarly, aboveground soluble carbohydrates showed associations with elevation.

**Fig 4.**
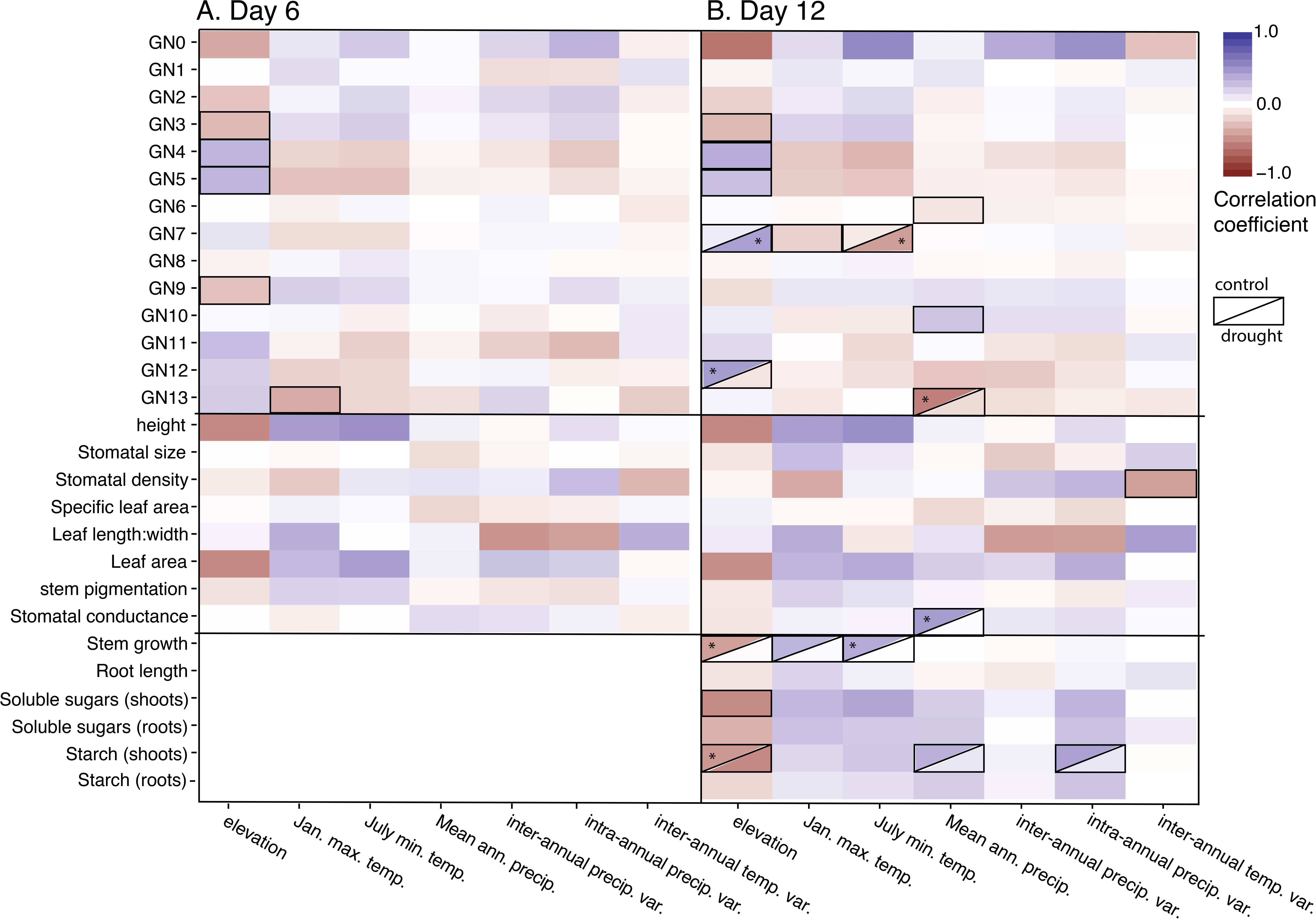
Heatmap of standardized regression coefficients for climate variables as predictors and traits and gene networks (GN) as response variables for measurements (A) on day 6 or (B) day 12. For variables with significant interaction effects, each treatment was analyzed separately to estimate slopes and test their significance within treatments. Solid boxes indicate a significant association and the box is divided in cases where there is a significant treatment and climate effect, indicating that the climate association was significant only in one of the treatments. The upper half of the box represents the watered treatment and the lower half represents the dry treatment. Stars indicate significant correlations for either of the two treatments.

We tested for climate by treatment interaction which revealed if associations between traits and the climate are significant for only one of the treatments. We found that gene expression patterns for GN7 at day 12 show a positive correlation with elevation (and minimum winter temperature) only for plants under drought conditions. Contrastingly, the correlations between GN12 and elevation, and GN13 and mean annual precipitation are significant only for watered plants. Similarly, for stem growth and aboveground starch, the correlations with elevation are significant only for watered plants, and stomatal conductance with mean annual precipitation only under watered conditions.

#### Plasticity differentiation is associated with climate

We tested if plasticity in traits and gene expression reflects signatures of climatic effects. At day 6, plasticity in GN5 (related to cytoskeleton organization) is the only gene network that show a correlation with a climatic variable: low elevation populations showed higher plasticity in gene expression patterns of this gene network (**Table 4**). We also observe higher plasticity in stomatal density in wetter sites. At day 12, we found that plasticity in GN10 (related to translation) and GN13 (linked to homeostasis) covaries with mean annual precipitation. For GN10, populations from both the driest and the wettest sites show higher plasticity, and populations from wetter sites show higher plasticity for GN13 (**Table 4**, **Figure 5D**). Additionally, plasticity in stem growth correlates with both elevation and summer maximum temperature. High plasticity in growth is observed in populations from low elevation and warmer sites (**Figure 5A**). Similar to GN13, plasticity in leaf area shows correlations with mean annual precipitation with higher plasticity in populations from wetter sites (**Figure 5B**). Pigment accumulation shows correlations with temperature heterogeneity (temperature range within years) (**Figure 5C**) and maximum temperature. However, pigment-related associations are driven by a single population in Cederberg, the most northern population we studied. We should also note that this trend is driven by the direction of plasticity in this population for pigment content, which is opposite to what we observe in all the other populations:, plants from the Cederberg population have higher pigment under drought and lower pigments when watered (Figure S3).

**Table 4.** Significant effects from mixed models testing associations between climate variables and plasticity measures, for traits and eigengenes of gene networks (GN) on day 6 and day 12.

| Day | Plasticity measure | Climate variable | p-value | Plasticity high in pops. from: |
| --- | --- | --- | --- | --- |
| Day 6 | GN5 | Elevation | 0.015 | lower elevation |
| Day 6 | Stomatal density | Mean annual precip. | 0.008 | wetter sites |
| Day 12 | GN10 | Mean annual precip. | 0.035 | wettest and driest sites |
| Day 12 | GN13 | Mean annual precip. | 0.021 | wetter sites |
| Day 12 | Growth | Elevation | 0.014 | lower elevation |
| Day 12 | Growth | Summer max. temp. | 0.016 | warmer sites |
| Day 12 | Leaf area | Mean annual precip. | 0.019 | wetter sites |
| Day 12 | Pigment accumulation | Temperature range | 0.039 | higher heterogeneity |
| Day 12 | Pigment accumulation | Summer max. temp. | 0.041 | coolest and warmest sites |

**Fig 5.**
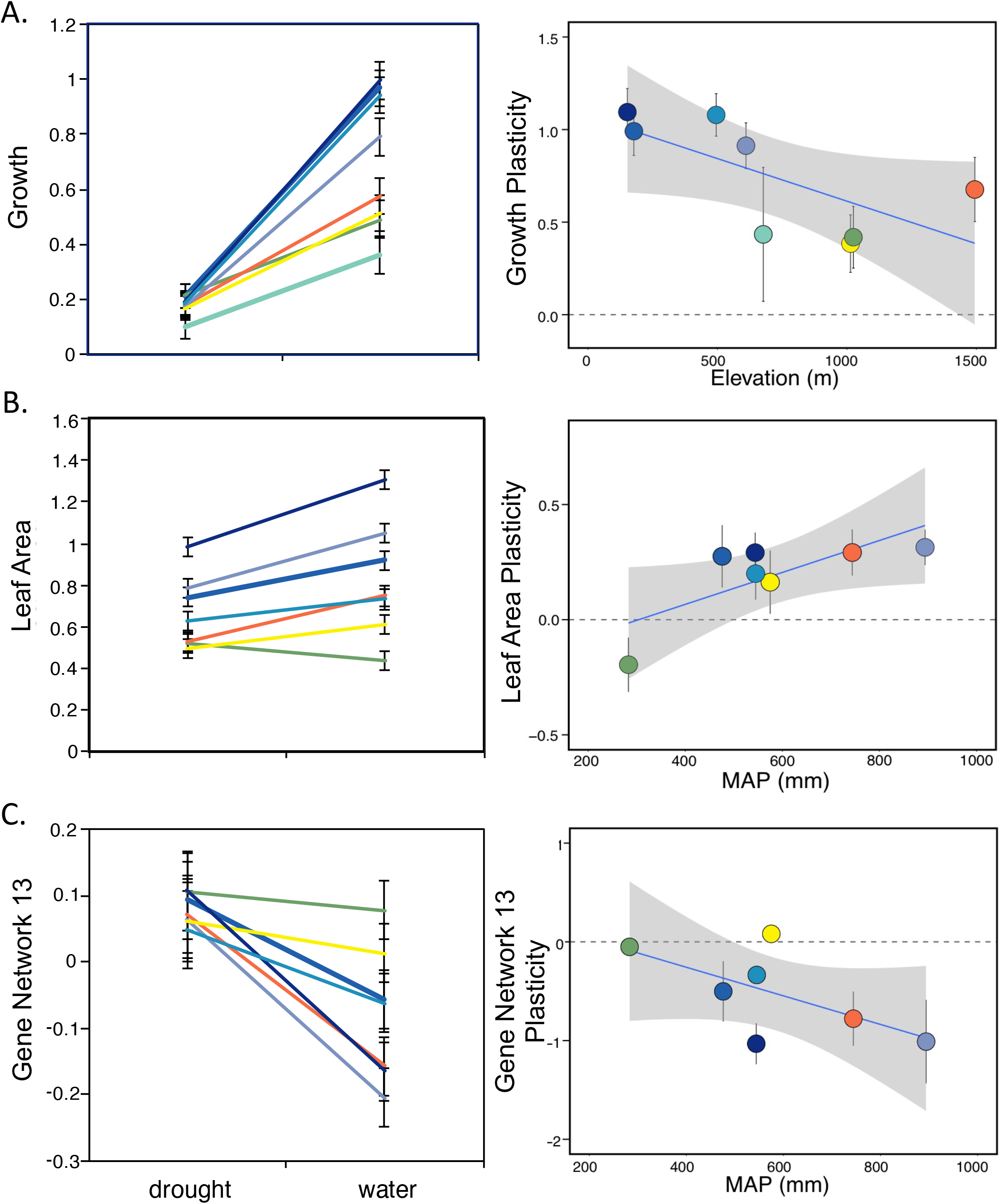
Response curves for the studied 8 populations under well-watered and drought conditions and the associations between plasticity and climate variables for growth (A), leaf area (B) and gene network 13 (C).

## Discussion

Local adaptation can lead to trait divergence among populations along environmental gradients. Plasticity in traits, as any other phenotypic character, can also be under selection, creating matching patterns between plasticity differentiation and the source site climatic variation among populations. In this study, we show that functional traits and gene expression patterns, and their plasticity are associated with source-site climate in *Protea repens*, suggesting local adaptation in both trait means and their plasticity.

### Drought imposes a complex stress regulating many gene expression networks

Drought has been a strong driver of plant evolution in the Cape Floristic Region and other Mediterranean climate zones (Allsopp, Colville, and Verboom 2014). When water availability is limited, as it typically is for several months during the summer in most of the region, plants experience a suite of complex stresses (West, Dawson, and February 2012; Ramachandra Reddy, Chaitanya, and Vivekanandan 2004) reflected in regulation of various pathways including ABA-dependent pathways, carbohydrate metabolism, ROS scavenging and photosynthesis under drought (Heschel et al. 2014; Miller et al. 2010). Accordingly, we observe differential gene expression in 10 out of 13 gene networks under drought conditions. For example, the genes with the highest gene network membership in GN5 (related to cytoskeleton organization) are downregulated under drought in all the populations, and this gene network also shows correlations with height, leaf area and aboveground carbohydrate content. Cytoskeleton organization related pathways were also shown to be highly hampered during drought stress in Arabidopsis (Bhaskara et al. 2017), linking this process to reduced growth. A general pattern similar to that of GN5 is seen in other gene networks important for growth (such as GN1 related to cell wall organization and other carbohydrate metabolism related gene networks). This “shut-off” of growth also explains the shorter stems and smaller carbohydrate content aboveground under 12 days of drought for all populations. Additionally, the genes that were consistently up-regulated among all the populations were genes involved in stress responses. Together these results indicate a trade-off in which resources are channeled from growth to stress acclimation. A similar trade-off has been observed across many plant species for herbivory (Kempel et al. 2011; Züst, Rasmann, and Agrawal 2015), cold stress (Loehle 1998) and drought (Haugen et al. 2008). We also detected reduced stomatal conductance in all populations in the drought treatment. This indicates a potential drop in photosynthesis, which might further contribute to the energy deficit hampering growth. The only gene network, GN6 (enriched for photorespiration related genes) that shows a correlation with conductance at the later stage of the stress is also correlated with leaf area and carbohydrate content. Interestingly, we also find a correlation between this gene network and mean annual precipitation suggesting local adaptation to water availability via regulation of carbohydrate accumulation.

### Associations with the climate are context dependent

When there are fitness consequences of a trait that is induced conditionally in response to environmental stressors, with no fitness costs/benefits when those stressors are absent, genetic variation that controls trait plasticity will be under selection only under the inducing stressful environment (Van Dyken and Wade 2010; Ledón-Rettig et al. 2014). Strikingly, we observed climatic correlations in gene expression and traits that only exist in either one of the treatments (**Figure 4**). Gene network 7, for example, shows a correlation with elevation and winter minimum temperature only when the plants are under drought stress. Gene ontology enrichments in this module point to regulation by high light and temperature stress. A similar gene network in our previous study in a common garden with more mature plants (18 months old) was also enriched for GO terms related to high light intensity and temperature, and this module showed correlations with stem diameter and stomatal density, but surprisingly not with any climate variables (Akman et al. 2016b). GN7 in this study also shows significant correlations with height at both timepoints and SLA in the earlier timepoint (**Table S6**). The top members of this gene network with Arabidopsis homology are down-regulated in populations coming from higher elevation, one of these genes is 4CL3 (AT1G65060) involved in last step of phenylpropanoid pathway - the upstream pathway anthocyanins are derived from. We also found that plants from warmer sites (higher maximum summer temperature) have higher pigment accumulation (p-value =0.029, fdr-corrected value = 0.106). Previous studies showed a negative effect of heat stress on plant growth (Olszyk et al. 1998; Hasanuzzaman et al. 2013; Bita and Gerats 2013), and anthocyanin accumulation as a protective mechanism against heat, cold, oxidative stress and UV radiation (Hatier and Gould 2009; Steyn et al. 2009; Zeng et al. 2010; Wang et al. 2013). The higher amount of anthocyanins accumulated in populations originating from warmer sites in this study might enable a better protection against stressors such as reactive oxygen species, and thus lead to taller stature. In a parallel finding, under drought conditions, differentially expressed genes among populations were enriched for only oxidative stress related genes, suggesting selection on expression of these genes along the climate gradient.

Contrastingly, GN13 (homeostasis related; the best representative gene showing homology with an Arabidopsis heat shock protein-AT5G59720) shows a correlation with mean annual precipitation only in watered conditions. Heat shock proteins are shown to be involved in responses to stresses such as heat, cold, UV radiation, salt and drought (Chen, Feder, and Kang 2018). Under control conditions, plants from wetter source sites express genes in this network at lower levels compared to populations from drier sites, and show a bigger up-regulation under drought, eventually reaching similar levels of expression for all populations. This gene network also shows correlations with leaf area, SLA, stomatal pore size and aboveground carbohydrate content. In plants from drier environments, higher constitutive expression of these genes might be crucial for a prompt response to drought via regulation of stomatal pore size, leaf size and thickness. However, plants from populations that rarely experience drought might not benefit from constitutive expression of these genes, creating a mismatch between costs and benefits.

### Minimum temperature and elevation show widespread associations with trends in gene expression

Previously, we showed that functional traits and gene expression patterns as measured in a common garden with older plants (1.5 year old plants vs 2 months-old plants in this study) correlated with source site climate (Akman *et al*., 2016). Although using fewer populations in this study, we observed similar patterns in those correlations. For example, in both studies we observe associations between gene expression networks and minimum winter temperature, elevation and mean annual precipitation (note that minimum winter temperature and elevation are highly correlated: Pearson’s R = −0.91, **Table S2)**. Taken together, these variables have several associations with gene network variation, as we also found in the previous study. Although *Protea* populations throughout the CFR experience drought at least occasionally, the drought metrics used in this study are not as strong predictors of trait variation as temperature/elevation is under the tested conditions (drought and control in a greenhouse and a common garden). Since gene expression networks include all major groupings of coexpressed genes, they reflect an unbiased pattern in plant responses, and in this case, this pattern indicates fairly widespread variation in gene expression across temperature/elevation gradients.

Why was temperature/elevation associated with population differences in gene expression more often than precipitation? We can’t say conclusively, but note that populations in the CFR experience drought periodically, and although some of the populations we studied experienced a decrease in precipitation over the past 115 years, this effect is not as strong as the increase in temperature over the same period (**Figure 6**, (Kruger and Shongwe 2004; Mason and Jury 1997). Both historically and during recent warming, temperature regimes potentially impose a more continuous selection force than precipitation patterns. Precipitation has been more variable historically and recent climate change trends are variable across the region, and this variability might make it harder to detect patterns of selection in response to precipitation changes across climate gradients even if measurements are performed under drought conditions as in our experiments. These results also line up with a broader trend: a meta-analysis of 21 plant traits from 25354 species concluded that temperature is a stronger driver of plant trait evolution compared to precipitation (Moles et al. 2014).

**Fig 6.**
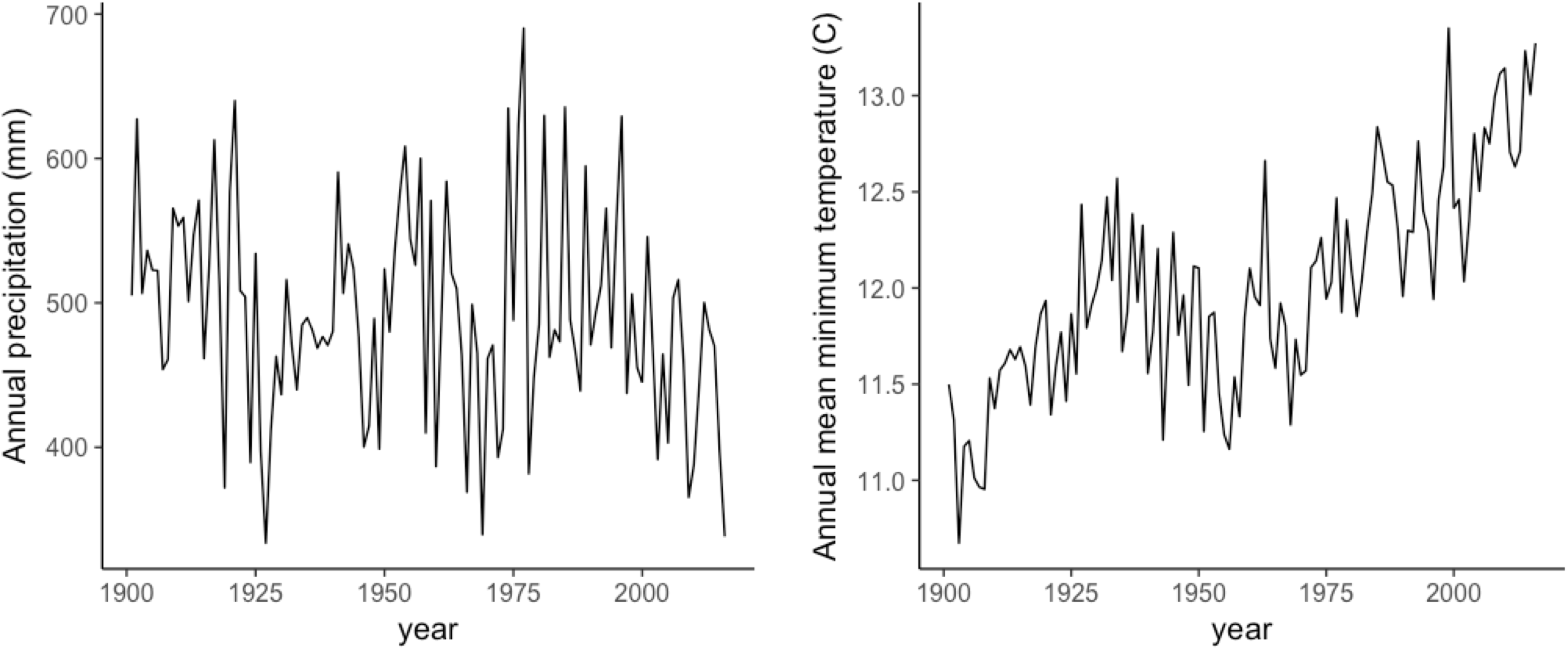
Mean annual precipitation (A) and mean annual temperature (B) averaged across the 8 population sites between 1901-2016.

### Carbohydrate reserves play a role in survival

Surprisingly, we found a positive correlation between survival under drought and precipitation; plants from wetter sites survived longer under drought. Populations from wetter sites also showed higher stomatal conductance and shoot starch content, indicating higher photosynthetic rates. The fact that populations from wetter sites survived better under drought is counterintuitive. However, in a sudden drought onset, like the one imposed in this experiment, carbohydrate reserves might be a decisive factor in survival, especially in earlier stages of development, and thus if plants from wetter sites have built up larger reserves, they may survive longer (O’Brien et al. 2014).

### Both climatic heterogeneity and favorable environmental conditions promote plasticity evolution

Here, we also tested if plasticity is favored and thus selected for in heterogeneous and/or milder climatic conditions. A heterogeneous environment in theory should promote plasticity evolution as this can allow for a better match between the phenotype and the changing environment, potentially surpassing cost of plasticity. Similarly, in mild environments, plasticity can arise through a relaxed selection in which a mismatch between the phenotype and the climate does not impose strong fitness costs. We found evidence for both of these patterns for the functional traits and the gene expression patterns. However, in this study, we have more evidence for growth plasticity evolving under milder conditions than heterogeneous environments, i.e. higher plasticity for growth and GN5 (cytoskeleton organization) in populations from lower elevation/warmer sites (representing milder climates). Although heat stress can be detrimental, in the CFR cold stress may impose a stronger selective pressure for growth (Wilson, Latimer, and Silander 2015). In fact, in a study of 56 Proteaceae species looking at climate and plant trait correlations, Mitchell et al, showed the most strongly supported evolutionary correlations to be between winter minimum temperature (next to mean annual temperature and elevation) and plant height (Mitchell, Carlson, and Holsinger 2018). Authors also note that this negative effect of elevation on plant height might be coupled potentially with shallower soils in the high elevation sites (Campbell and Werger 1988). These combined effects might create harsher environments which increase the cost of mismatch between phenotype and the environment, thus decreasing fitness of more plastic individuals. We also find higher plasticity in wetter sites for GN13 (homeostasis related) and leaf area, and again these sites represent milder climates in this drought-prone region. The only correlation we observe for heterogeneity is also related to temperature: plasticity in pigment accumulation increases with increasing heterogeneity. In environments where temperature change across seasons is high, levels of anthocyanins might effectively protect plants against abiotic stressors, thus promoting selection on higher plasticity.

To conclude, our results suggest that both phenotypic traits and gene expression, and their plasticity can be under selection along a climate gradient. However it is also important to note that without studying fitness consequences of the climate, our results do not prove that this variation is adaptive. We also showed that detecting associations between these traits and the environmental variables can be a challenging task as these associations are environment-dependent, and thus can be detected only when they are measured under certain conditions. Additionally, we found that trait variation can be explained better with temperature related-variables rather than precipitation related variables we chose. This could arise from recent temperature increase associated with climate change that can potentially impose a stronger and more continuous selection pressure compared to the more variable changes in precipitation. This study also provides insights into the drivers of plasticity evolution: we found more evidence for higher plasticity in mild environments compared heterogeneous environments. Although our results demonstrate population differentiation in traits and their plasticity and their links to climate, we should also note that we found only 24/350 trait/gene expression and climate associations to be significant together with 9/33 for plasticity. This relatively low number could either be due to the fact that the climate variables we chose cannot fully represent the source site environments, that our experimental design does not capture the genetic variation fully, or that drift has affected population differentiation. In this regard, future work with long-term transplant experiments in several common gardens and collection of genomics/transcriptomics data would enable heritability and Fst/Qst calculations, and coupling these with fitness measurements would conclusively uncover patterns of local adaptation and the underlying genetic/gene expression mechanisms in this vulnerable species in the CFR.

## Acknowledgements

We thank Chris Adams, Megan Varvaro and Sarah Soorya for their assistance with plant care and the drought experiments, Tony Rebelo, Guy Midgley and Louise Nurrish for their assistance with seed collection. We thank Jessica Orozco and the Zwieniecki Lab for their contribution in the carbohydrate analysis. We also thank the Blackman Lab and Carl Schlichting for their suggestions on an earlier version of the manuscript. Finally, we thank the reviewers for providing detailed constructive suggestions that increased the quality of our manuscript. This work was supported by Dimensions of Biodiversity awarded to Carl Schlichting, Cynthia S. Jones and Kent E. Holsinger (award no: 1046328) and Andrew M. Latimer (award no: 1045985).

## Data Accessibility

All sequencing data will be made available at NCBI SRA. Bioinformatics and R scripts will be made available upon acceptance of the manuscript on Akman’s github. Climate data, phenotypic measurement and gene expression counts will be made available in Dryad.

## Author contributions

Akman, Carlson and Latimer designed the research. Akman and Carlson performed the dry down experiments. Akman prepared sequencing libraries and performed gene expression analyses. Akman, Carlson and Latimer performed climate association analyses and wrote the manuscript.

## SUPPLEMENTARY INFORMATION

Table S1. Population source site climatic information

Table S2. Correlations between climatic variables

Table S3. Maternal lines used in the experiment, with sample sizes for each maternal line in the two treatments (drought and watered)

Table S4. WGCNA GO categories and highest gene network membership genes

Table S5. Up and down-regulated genes for all the populations under drought

Table S6. Gene network and trait correlations

Table S7. Results of linear mixed effect models for climate, treatment and climate x treatment effects for traits.

Table S8. Results of linear mixed effect models for climate, treatment and climate x treatment effects for gene networks.

Figure S1. Sampling scheme

Figure S2. WGCNA sample clustering

Figure S3. Functional traits response curves

Figure S4. Gene network response curves

Figure S5. Performance traits response curves

Figure S6. MDS plot with all samples for day 6 and day 12

Supplementary Materials and Methods

