## Supplementary Information for "Climate explains population divergence in drought-induced plasticity of functional traits and gene expression in a South African *Protea*"

***Supplementary Materials and Methods***

**Transcriptome sampling**

A smaller subset of plants compared to functional trait measurements had one leaf per plant harvested for whole transcriptome sequencing and gene expression analysis (n=120 plants total, or 4 per population per treatment per day, except no ALC on day 12 due to insufficient number of individuals because of lower germination in this population). Plants with leaves removed on day 6 were not sampled for transcriptomics on day 12, to avoid possible shifts in gene expression caused by leaf removal. As a consequence, gene expression and most traits were measured in one set of plants on day 6 and in a separate set on day 12. Transcriptome sampling took place prior to any other measurements on a given day and consisted of collecting one leaf for RNA extraction per plant (the 2nd or 3rd youngest leaf from the tip) into a 1 mL tube and snap-freezing immediately in liquid nitrogen. These samples were stored at -80°C until library preparation. All sampling for transcriptome sequencing was done between 10 AM and 11 AM, and functional trait measurements were done between 10AM and 2PM each day. Both transcriptomics and other trait measurements were done in a random order. On day 12, we harvested a subset of plants for root length and carbohydrate storage measurements (n=9 per population including the 4 transcriptome plants and 5 additional plants). After completing all trait and transcriptome sampling, we cut plants at the root- shoot junction and placed aboveground tissues in a 10 mL tube and snap-froze the samples in liquid nitrogen for future carbohydrate analyses. Belowground tissues were gently removed from the soil, rinsed clean and stored in a separate tube and also snap-frozen. For these 9 plants per population, we also measured the length of the longest root. Tubes containing plant tissues were stored in a -80°C freezer for ~1 month, then freeze-dried for biomass and carbohydrate analyses. Dried tissues for above- and belowground material was measured and ground. Ten to hundred and fifty mg of ground material was used for carbohydrate analyses as described in (Akman *et al.* 2012). Briefly, carbohydrates and starch were extracted with 70% methanol solution. The extract was used for soluble carbohydrate content measurements and the pellet including starch was treated enzymatically with α-amylase and amyloglucosidase resulting in hexose sugars.

Soluble carbohydrates and hexose sugar concentration was measured with a modified version of anthrone method using fixed glucose standards. We also measured biomass but biomass and carbohydrate content were highly correlated, and thus only carbohydrate content results are shown (dry mass and soluble sugars Pearson's R = 0.90 and 0.94; starch Pearson's R = 0.64 and 0.81 for belowground and aboveground, respectively).

**Transcriptome library construction and sequencing**

Transcriptome sequencing samples were homogenized in a Beadbeater-96 (Biospec Inc, Bartlesville, OK, USA) with 2.3 mm Chrome-Steel Beads (Biospec Inc, Bartlesville, OK, USA). Lysis/binding buffer (Life Technologies, Grand Island, NY, USA, with addition of %3 PVP-40) was added and the solution was homogenized again for 2 min. Tubes were transferred to a 60°C water bath for 30 min with occasional shaking every 10 min. The solution was centrifuged for 3 min at 12 000 x g and the supernatant was used for mRNA isolation and sequencing library preparation by the procedures explained in Kumar et al. Sequencing libraries were pooled and quality control of the libraries were done with Bioanalyzer (Agilent Technologies Inc., Santa Clara, CA, USA). Before pooling, concentration of each library was measured in a plate reader and appropriate amounts were pooled in order to balance reads obtained from each library from the pooled samples. A total of 120 samples were sequenced; 60 in each of 2 Illumina Hiseq 2500 lanes, yielding 756 million single-end reads (100 bp long). A randomized block design was used for sequencing: two samples per population per treatment per time-point were included in each lane.

**Survival measurements**

After all trait and gene expression data had been collected, the remaining plants in the drought treatment (those that had not been harvested at the end of the experiment) were maintained in the greenhouse under drought stress with regular watering. Plants were checked weekly to score survival and days until mortality was recorded.

**Associations between traits, gene expression and their plasticity and survival**

None of the plants in the control/watered treatment died during our experiments, however we observed mortality under drought conditions. Survival times ranged from 4 to 47 days after imposition of the drought treatment. We used linear mixed models to test for associations between trait means and plasticity levels and survival time. We tested for association with survival by regressing the length of time until death on the trait mean or plasticity value, with population as a random effect. We compared models that included that fixed effect of the trait to a null model that included only the population random effect, using chi-square tests. We also tested for associations between source site climate and survival by first averaging up the survival values to the population level, since we have only environmental data value per population, then using linear regression to relate survival to environmental factors.
