## Supplementary material for "Climate explains population divergence in drought-induced plasticity of functional traits and gene expression in a South African *Protea*": Table S1 and S2

**Table S1.** GPS coordinates and climate characteristics of the 8 seed source populations of *Protea repens*.
Climate data are from Schulze 2007 except for rainfall seasonality and inter-annual rainfall CV, which both came from Schulze 1997 (respectively called: RAINFALL CONCENTRATION: Based on Markham's Technique, and INTER‐ANNUAL COEFFICIENT OF VARIATION (%) OF PRECIPITATION).

| **Seed**  **source** | **Lat.** | **Long.** | **Elev.**  **(m)** | **Mean annual rainfall** | **Jan. max.**  **temp** | **July min.**  **temp.** | **Inter-annual rainfall CV** | **Rainfall seasonality (within year)** | **Annual max. – min. temp.** |
| --- | --- | --- | --- | --- | --- | --- | --- | --- | --- |
| A | -33.2408 | 26.0984 | 677 | 429 | 27 | 4.7 | 32.25 | 18 | 22.3 |
| B | -34.5459 | 20.0386 | 177 | 476 | 27 | 6.6 | 34.8 | 25 | 20.4 |
| C | -32.4064 | 19.1068 | 1014 | 575 | 19.9 | 2.8 | 32.1 | 56 | 17.1 |
| F | -33.3654 | 19.2760 | 610 | 894 | 27.3 | 1.9 | 33.2 | 53 | 25.4 |
| G | -33.9679 | 21.2198 | 495 | 545 | 30.2 | 5 | 23.9 | 5 | 25.2 |
| R | -33.5160 | 18.5511 | 152 | 544 | 29.5 | 5.9 | 27.4 | 52 | 23.6 |
| S | -33.3495 | 22.0459 | 1494 | 744 | 25.8 | -0.1 | 29.9 | 5 | 25.9 |
| V | -33.4934 | 23.6336 | 1025 | 283 | 27.1 | 1.1 | 23.3 | 13 | 26 |

**Table S2.** Pearson’s R correlations among climate variables. Ordered as strongest to weakest associations.

| **Variable 1** | **Variable 2** | **Pearson’s R** |
| --- | --- | --- |
| Elevation | July min.temp. | -0.91 |
| Jan. max.temp | Annual max. – min. temp. | 0.70 |
| Inter-annual rainfall CV | Annual max. – min. temp. | -0.56 |
| Elevation | Jan. max.temp | -0.52 |
| Rainfall seasonality | Annual max. – min. temp. | -0.49 |
| Inter-annual rainfall CV | Annual max. – min. temp. | -0.44 |
| Inter-annual rainfall CV | Rainfall seasonality | 0.43 |
| Inter-annual rainfall CV | Mean ann. rainfall | 0.42 |
| July min.temp. | temp range | -0.40 |
| Rainfall seasonality | Jan. max.temp | -0.37 |
| Jan. max.temp | July min.temp. | 0.37 |
| Rainfall seasonality | Mean ann. rainfall | 0.35 |
| Rainfall seasonality | Elevation | -0.33 |
| Mean ann. rainfall | July min.temp. | -0.32 |
| Elevation | Annual max. – min. temp. | 0.18 |
| Mean ann. rainfall | Annual max. – min. temp. | 0.16 |
| Rainfall seasonality | July min.temp. | 0.16 |
| Inter-annual rainfall CV | July min.temp. | 0.16 |
| Elevation | Mean ann. rainfall | 0.15 |
| Inter-annual rainfall CV | Elevation | -0.13 |
| Mean ann. rainfall | Jan. max.temp | -0.08 |
