## Supplementary material for "Climate explains population divergence in drought-induced plasticity of functional traits and gene expression in a South African *Protea*": Table S3

**Table S3. Maternal lines used in the experiment, with the sample sizes for each maternal line in the two treatments (drought and watered).**

| **Population** | **Maternal Line** | **Drought** | **Watered** |
| --- | --- | --- | --- |
| A | A.34 | 2 | 2 |
| A | A.35 | 0 | 1 |
| A | A.38 | 1 | 1 |
| A | A.43 | 2 | 0 |
| A | A.44 | 1 | 1 |
| B | B.15 | 1 | 1 |
| B | B.19 | 2 | 3 |
| B | B.22 | 1 | 1 |
| B | B.24 | 1 | 1 |
| B | B.25 | 3 | 3 |
| B | B.34 | 1 | 1 |
| B | B.35 | 2 | 1 |
| B | B.37 | 2 | 3 |
| B | B.39 | 1 | 1 |
| B | B.42 | 1 | 1 |
| C | C.10 | 1 | 1 |
| C | C.12 | 1 | 0 |
| C | C.15 | 2 | 2 |
| C | C.18 | 1 | 1 |
| C | C.19 | 0 | 1 |
| C | C.20 | 2 | 1 |
| C | C.22 | 1 | 1 |
| C | C.23 | 0 | 1 |
| C | C.25 | 2 | 5 |
| C | C.26 | 1 | 1 |
| C | C.27 | 1 | 0 |
| C | C.40 | 1 | 1 |
| F | F.1 | 3 | 3 |
| F | F.10 | 3 | 3 |
| F | F.23 | 0 | 1 |
| F | F.26 | 3 | 3 |
| F | F.28 | 3 | 3 |
| F | F.30 | 3 | 3 |
| F | F.36 | 3 | 4 |
| F | F.4 | 1 | 0 |
| F | F.40 | 3 | 3 |
| F | F.41 | 1 | 0 |
| F | F.5 | 2 | 3 |
| G | G.12 | 3 | 2 |
| G | G.26 | 3 | 3 |
| G | G.30 | 2 | 3 |
| G | G.33 | 4 | 4 |
| G | G.35 | 4 | 6 |
| G | G.36 | 3 | 2 |
| G | G.37 | 0 | 1 |
| G | G.38 | 3 | 3 |
| G | G.39 | 3 | 3 |
| G | G.7 | 2 | 3 |
| R | R.16 | 1 | 1 |
| R | R.17 | 2 | 0 |
| R | R.2 | 2 | 3 |
| R | R.23 | 2 | 2 |
| R | R.24 | 1 | 1 |
| R | R.25 | 2 | 3 |
| R | R.29 | 2 | 0 |
| R | R.3 | 1 | 2 |
| R | R.31 | 2 | 4 |
| R | R.34 | 3 | 2 |
| R | R.4 | 3 | 2 |
| R | R.41 | 2 | 3 |
| R | R.5 | 3 | 3 |
| S | S.10 | 2 | 2 |
| S | S.11 | 4 | 5 |
| S | S.15 | 2 | 0 |
| S | S.2 | 1 | 0 |
| S | S.22 | 2 | 1 |
| S | S.26 | 1 | 1 |
| S | S.30 | 2 | 4 |
| S | S.37 | 3 | 3 |
| S | S.41 | 3 | 3 |
| S | S.42 | 2 | 2 |
| S | S.45 | 2 | 3 |
| V | V.16 | 1 | 2 |
| V | V.20 | 2 | 3 |
| V | V.26 | 2 | 2 |
| V | V.27 | 1 | 0 |
| V | V.3 | 1 | 1 |
| V | V.35 | 1 | 1 |
| V | V.37 | 3 | 3 |
| V | V.38 | 1 | 2 |
| V | V.41 | 1 | 1 |
| V | V.42 | 3 | 4 |
| V | V.45 | 1 | 2 |
