## Supplementary figures and images for "Climate explains population divergence in drought-induced plasticity of functional traits and gene expression in a South African *Protea*"

### Figure S1

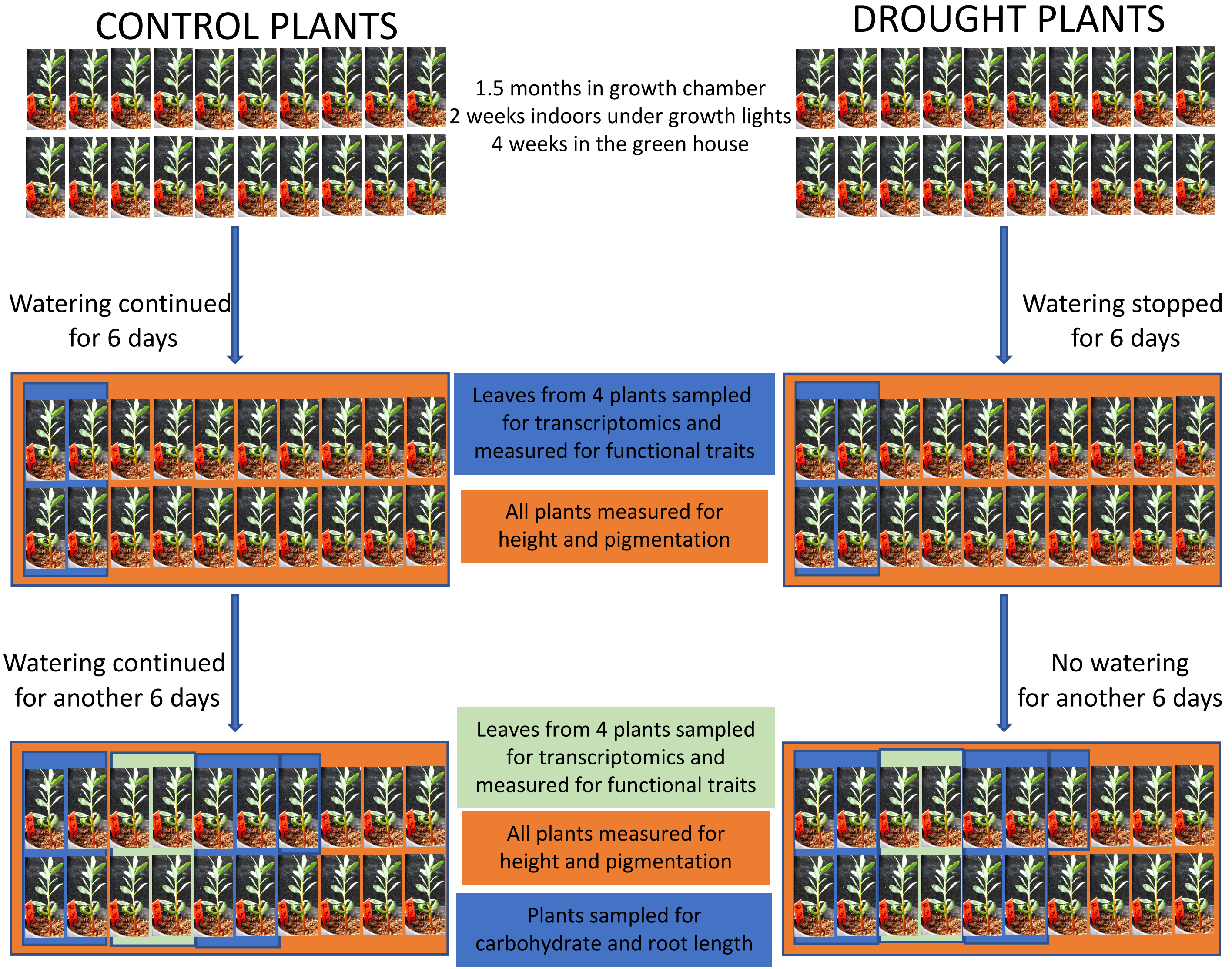


Figure S1. Sampling scheme

### Figure S2

#### Sample clustering on all genes in day1

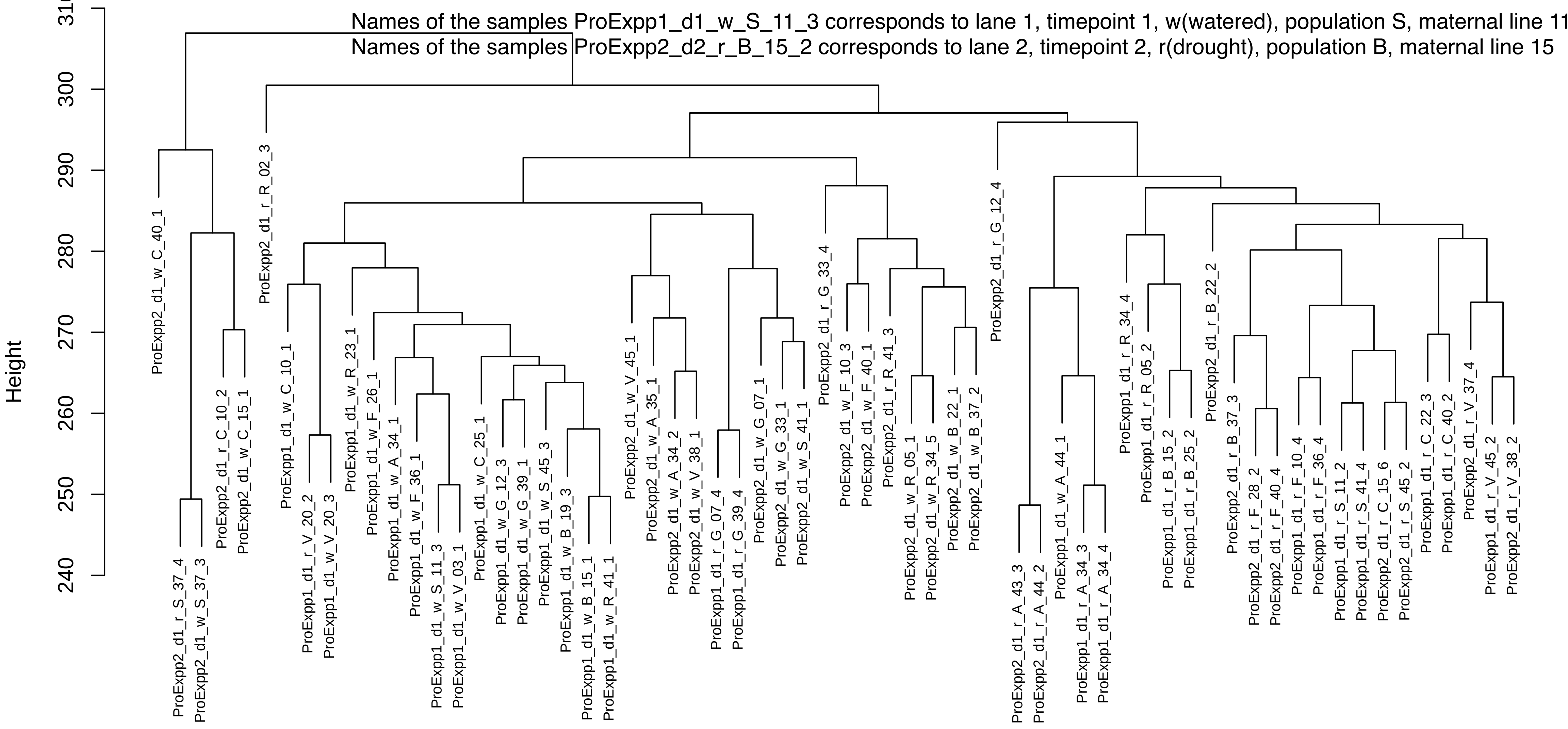


#### Sample clustering on all genes in day2

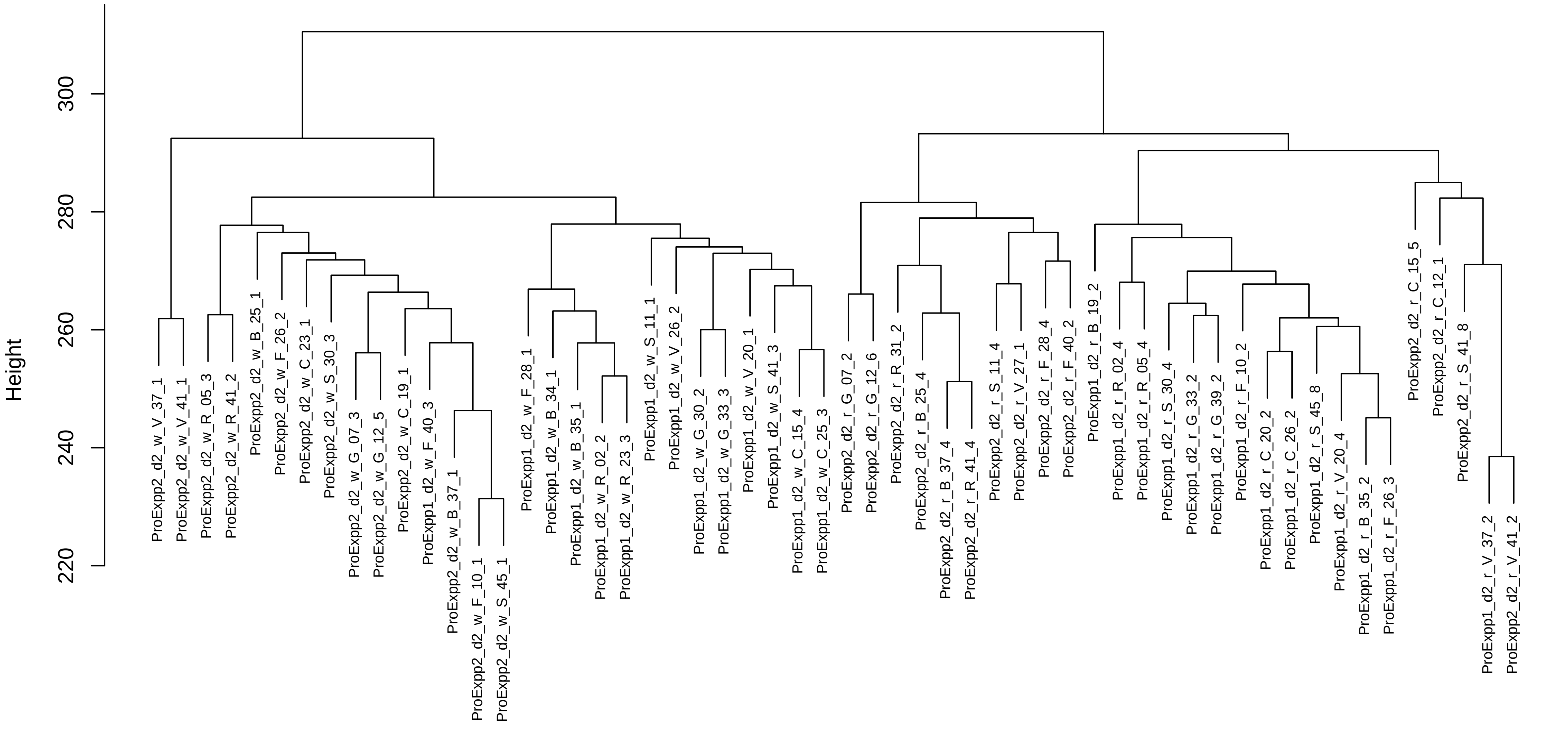

### Figure S3

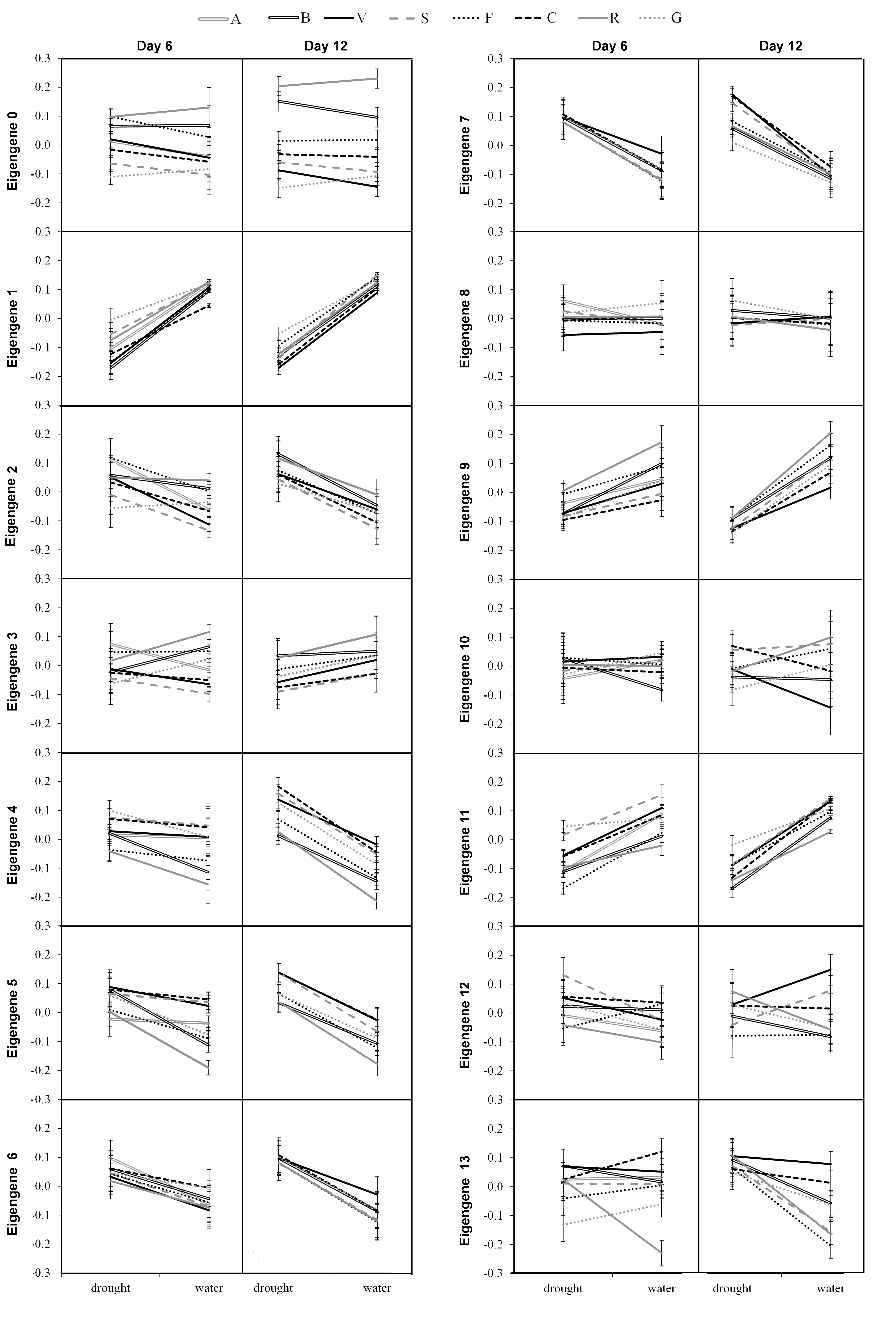

### Figure S4

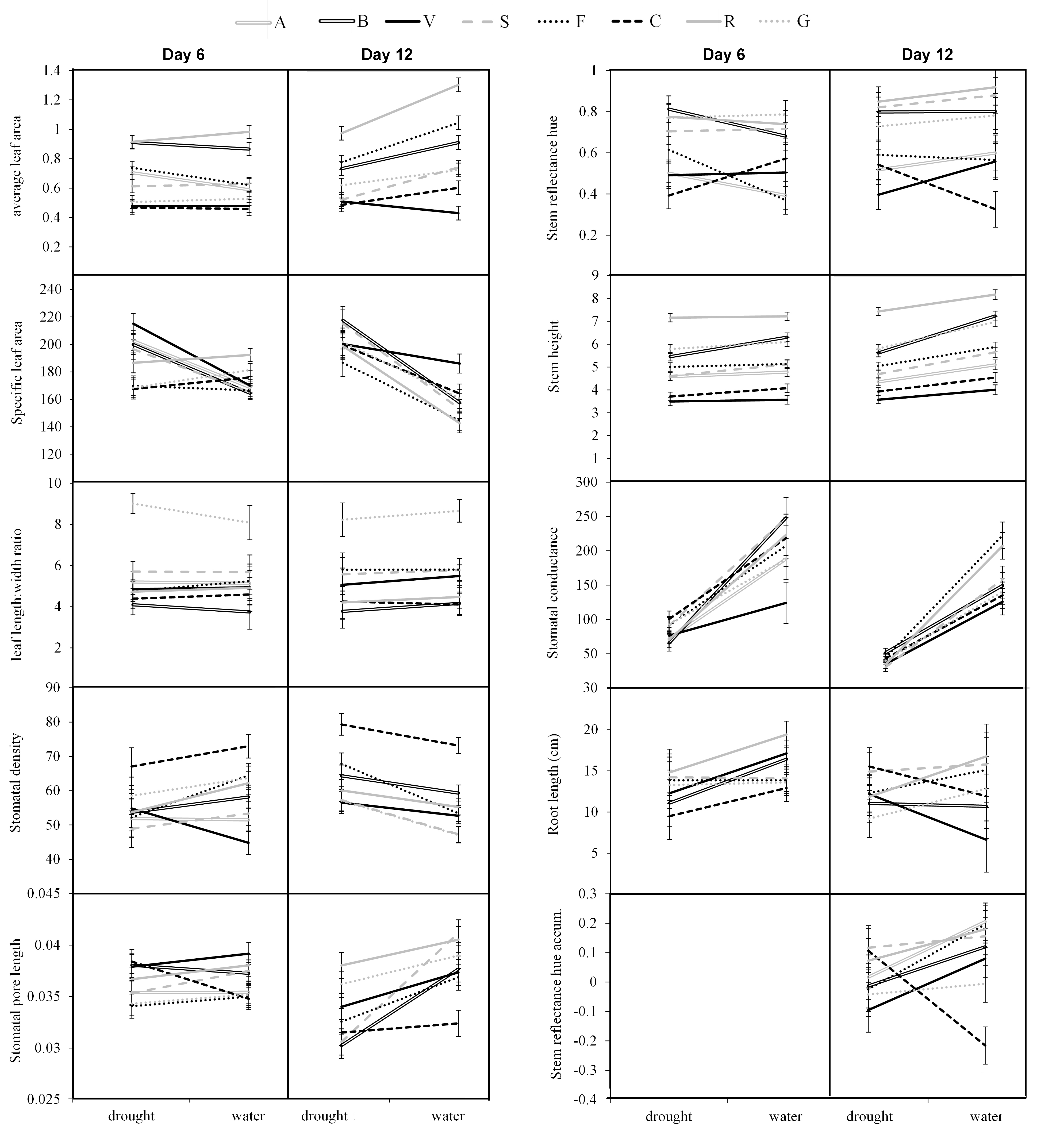

### Figure S5

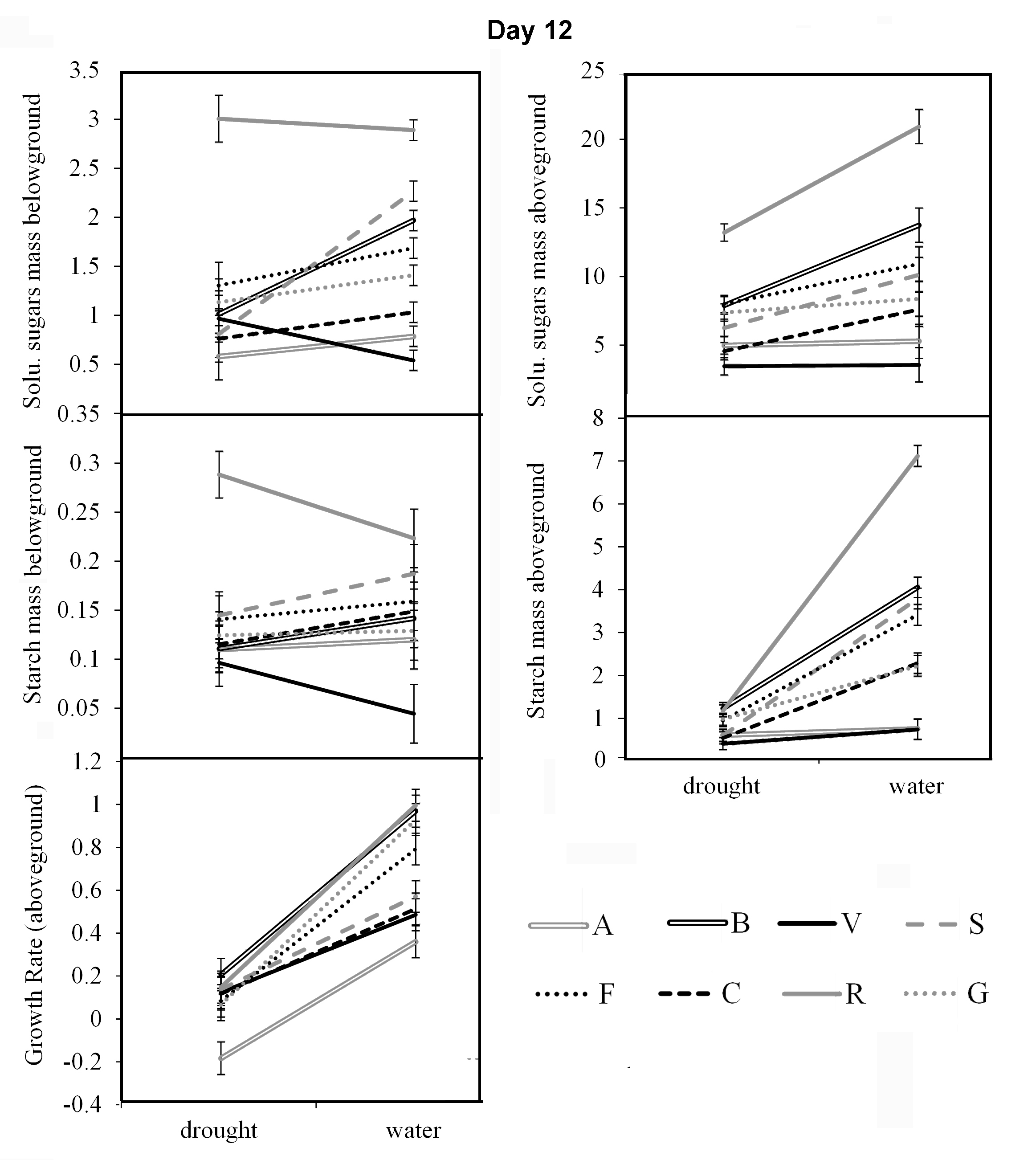

### Figure S6

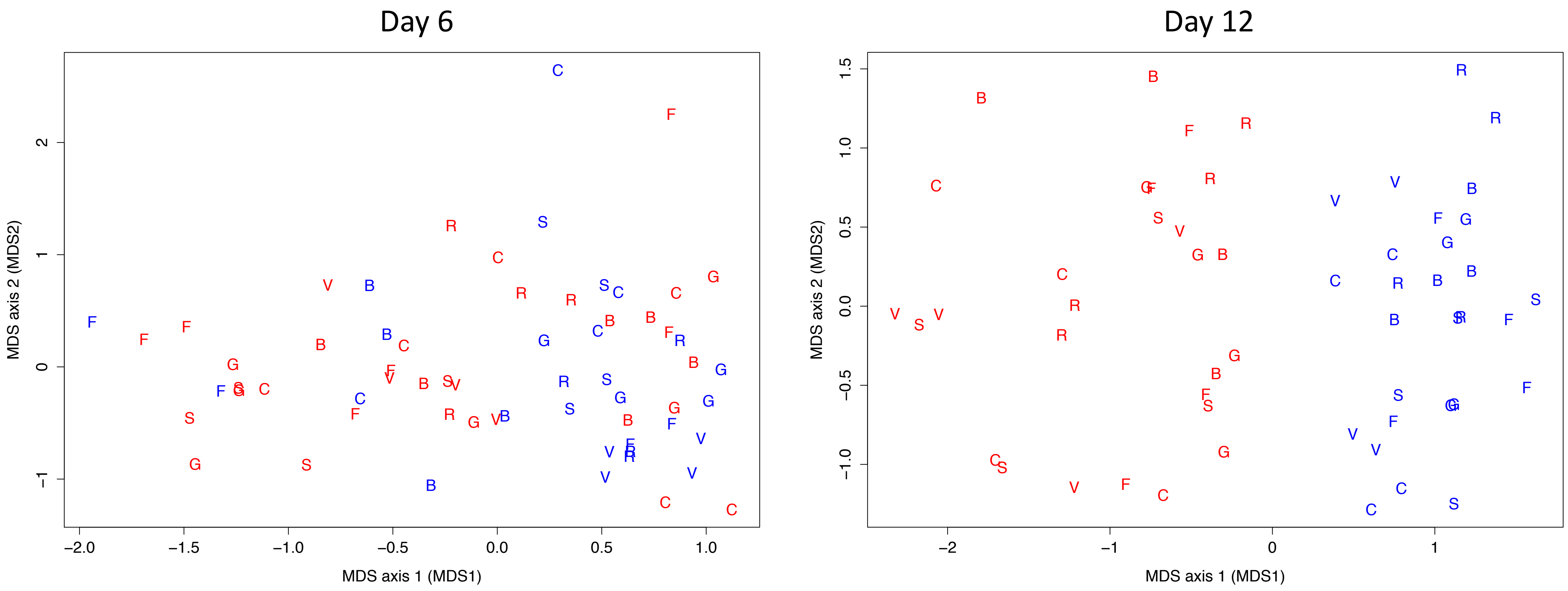


Blue control
Red drought
